# Generalizing Protein Fitness Prediction Across Distribution Shifts via Structural and Co-evolutionary Priors

**DOI:** 10.64898/2026.10.06.756934

**Authors:** Nopsinth Vithayapalert, Maytus Piriyajitakonkij, Francesca Grisoni

**Author notes:** Equal contribution (Nopsinth Vithayapalert, Maytus Piriyajitakonkij).

## Abstract

Protein fitness prediction is essential for protein engineering. Recent fitness-landscape benchmarks such as FLIP2 have shown that methods performing well under conventional train–test splits can fail when the test set is shifted out of distribution (OOD), including mutations at unseen positions and variants from unseen protein backbones. In particular, current protein language models (pLMs) often suffer substantial performance degradation under such distribution shifts, limiting their practical use in protein engineering. We therefore develop a framework that augments sequence representations with structural and co-evolutionary information for fitness prediction. Specifically, we pair structure-aware pLMs with features pooled from an AlphaFold-style trunk applied to each sequence and its multiple sequence alignment (MSA), and use a lightweight prediction probe to map these frozen representations to fitness scores. Our framework consistently outperforms the best previously reported FLIP2 scores across several types of distribution shift. We then systematically study the contribution of each component in our framework to generalization. Structural priors yield the largest performance gains on held-out protein backbones, whereas trunk features improve generalization to held-out mutation positions. Rather than proposing a single best method, we characterize which modalities are effective for different protein families and distribution shifts, providing practical guidance for choosing representations according to the generalization regime. These findings support the development of more generalizable fitness predictors that can serve as effective reward functions or oracles for protein design toward desired functions.

## 1 Introduction

Characterizing protein properties is essential in protein engineering. A protein’s sequence determines its structure (Anfinsen, 1973; Dill & MacCallum, 2012) and biophysical characteristics (Whisstock & Lesk, 2003; Park et al., 2024), which in turn influence experimentally measured properties such as solubility, stability, binding affinity, and turnover (Dieckhaus et al., 2024; van der Flier et al., 2024). A *fitness landscape* maps each protein amino acid sequence to the value of a measured property of interest (‘protein fitness’). An accurate and robust fitness predictor can therefore serve as an oracle that guides the search toward unexplored proteins with desired properties. Building such a predictor is hindered by the limited availability of experimentally measured fitness labels. Self-supervised representation learning, such as protein language models (pLMs) (Bepler & Berger, 2019; Rao et al., 2021), offers one way to address this bottleneck. These models encode protein sequences into latent representations that can transfer to fitness prediction using few training labels, achieving strong performance across existing benchmarks (Notin et al., 2023; Dieckhaus et al., 2024; Dallago et al., 2021; Didi et al., 2026).

Several existing benchmarks have been established to evaluate fitness prediction, such as the FireProt (Bednar et al., 2015) database, which aggregates manually curated thermostability measurements for single-point mutants, and ProteinGym (Notin et al., 2023), which collects hundreds of standardized deep mutational scanning (DMS) assays covering diverse functional properties. Their protocols, however, do not test whether the predicted sequence-structure-function relationships are preserved under out-of-distribution (OOD) shifts. This aspect is particularly key for wet-lab protein engineering, where later rounds deliberately move away from the training data – probing untouched positions, or switching wild type altogether. A predictor is useful only if it survives that move. The recent FLIP2 benchmark (Didi et al., 2026) addresses this concern by introducing data splits that explicitly evaluate generalization across such OOD shifts. It shows that fine-tuned pLMs do not necessarily outperform simple baselines, such as zero-shot scoring or one-hot features. Similarly, Talpir & Fleishman (2026) suggest that pLM-based approaches may be approaching their performance ceiling, highlighting the potential of complementary biophysical and structural information. Moreover, pLMs trained to capture biophysical properties (e.g., residue fluctuations, secondary structure, and inter-residue dynamic interactions) can improve the zero-shot prediction of mutation effects on protein folding stability (Hou et al., 2026). These findings motivate a systematic investigation of how features from diverse modalities complement one another and contribute to the generalization of protein fitness prediction.

In this work, we ask what sequence alone is missing to stretch the boundaries of fitness prediction, and test two candidates: (i) *geometric priors*, via structural tokens from structure-aware pLMs, and (ii) *co-evolutionary context*, via trunk representations from a structure-prediction model conditioned on sequence and multiple sequence alignment (MSA). Specifically, we extract single and pair representations from a pLM and the Protenix-v2 trunk (Zhang et al., 2026) and summarize them into nine fixed-dimensional feature blocks. Three blocks are obtained by global pooling over residues or residue pairs and are therefore wild-type (WT) independent, while the remaining six pool representations at, or relative to, the mutated positions and are WT dependent. We then use a lightweight prediction probe to map these frozen representations to fitness scores.

Concretely, we show that augmenting frozen pLM representations with structural and coevolutionary features substantially improves over conventional pLM baselines on the shifts where they struggle most, without fine-tuning any encoder — with the two modalities helping in distinct regimes: structural priors when generalizing to unseen protein backbones, and trunk representations when extrapolating to unseen mutation positions.

We then analyze *why* these representations help. An MSA-Pairformer control (Akiyama et al., 2026), which sees the same alignment without a structural objective, recovers a substantial fraction of the trunk gain on AMYLASE, isolating what geometry contributes beyond co-evolution. Separating in-distribution from out-of-distribution performance further shows that trunk features yield substantially larger gains OOD than in-distribution (ID) on the AMYLASE position splits, indicating genuine positional extrapolation rather than a better fit to the training data. Finally, wild-type-dependent, mutation-indexed feature blocks improve within-protein extrapolation on AMYLASE, quantifying how much knowing *where* a variant differs from its wild type is worth.

Ultimately, our contributions are:

1. A lightweight fitness predictor that pools frozen structure-aware pLM embeddings with single and pair representations from an AlphaFold-style trunk, requiring no fine-tuning and improving over the best published FLIP2 results on most position and wild-type splits.
2. A decomposition of where that gain comes from, separating co-evolutionary from geometric signal via an MSA-only control, and informative from extrapolative features via a paired ID/OOD evaluation.

## 2 Related works

**Protein language models.** pLMs are pretrained on natural sequence patterns; their embeddings in principle encode the determinants of protein function. For example, ESM-2 (Lin et al., 2023),

Dayhoff (Yang et al., 2025), and CARP (Yang et al., 2024) scales autoregressive or masked language models (MLM) over UniRef, while ESM-C enhances the representation quality by expanding its pretraining set to include metagenomic samples. Beyond sequences, structure-aware pLMs consume backbone structures to incorporate geometric priors as discrete structural tokens alongside residue tokens. SaProt (Su et al., 2024) pairs each residue token with a Foldseek 3Di token describing its local tertiary environment. ProSST (Li et al., 2024) instead learns its own quantizer over local structural neighborhoods and couples the two streams through disentangled attention. DPLM-2 (Wang et al., 2025) treats sequence and structure symmetrically, modeling both token types under a single multi-modal diffusion language model that can generate in both directions.

**Sequence-structure representation.** Recent structure prediction models (Jumper et al., 2021; Zhang et al., 2026; team et al., 2024; Passaro et al., 2025) learn sequence-structure representations through the trunk module that couples the co-evolutionary signal in a sequence’s MSA with geometric priors, yielding single representations that capture per-residue features and pair representations that encode residue–residue interactions. Prior works have explored applying atom-level trunk embeddings for various downstream tasks such as protein-ligand affinity prediction (Passaro et al., 2025), or small molecules’ ADMET profiles (Jang et al., 2026). On protein representations, Lu et al. (2025) develop tokenized all-atom structure embeddings, called CHEAP embeddings, which serve as a compact joint encoding of structure and sequence. Closer to the setting of this work, (Cheng et al., 2023) adapts AlphaFold 2 trunk Evoformer single and pair representations together with structural features to predict missense variant effects proteome-wide. The trunk’s attention maps are also shown to capture the epistatic residue-residue dependencies underlying functional stability. However, AlphaMissense focuses on variant-effect prediction for single amino acid substitutions. No prior work has systematically studied whether the information carried by these representations transfers to fitness prediction under distribution shift.

**Protein Fitness Prediction.** Protein fitness prediction has been approached in both unsupervised and supervised settings. In the unsupervised setting, pLM likelihoods provide zero-shot scores that have been shown to correlate with experimentally measured variant effects. In the supervised setting, pre-trained pLM embeddings can instead serve as features for lightweight regression models. In particular, linear probes provide strong baselines and can outperform more flexible non-linear models when labelled data are limited or cover only a narrow region of the fitness landscape. Prediction performance is commonly benchmarked on FireProt (Bednar et al., 2015) and ProteinGym (Notin et al., 2023). Recent work (Talpir & Fleishman, 2026) further shows that neither zero-shot pLM likelihoods nor supervised transfer from pLM representations consistently outperform conventional approaches such as homology-based scoring. However, these benchmarks primarily evaluate prediction within individual protein landscapes and do not explicitly test generalization under the out-of-distribution shifts encountered in practical protein engineering.

## 3 Method

To predict fitness scores, we fit lightweight non-linear regression models on features extracted from frozen encoders. We draw on three sources of information:

- **Sequence.** pLM gives a per-residue sequence representation *h ∈* R*^N×d^*^pLM^ , where *N* is the number of residues and *d*_pLM_ the pLM embedding dimension.
- **Structure.** Structure-aware pLMs additionally consume discrete structural tokens describing each residue’s local geometric environment, for which we encode the wild-type structure with Foldseek (van Kempen et al., 2022).
- **Co-evolution and geometry.** We pass each sequence, wild-type and mutant alike, together with its multiple sequence alignment (MSA) through the Protenix-v2 trunk (Zhang et al., 2026), yielding a single representation *s ∈* R*^N×^*^384^ that captures per-residue features and a pair representation *z ∈* R*^N×N×^*^256^ that encodes residue-residue interactions (Figure 1).

**Figure 1:**
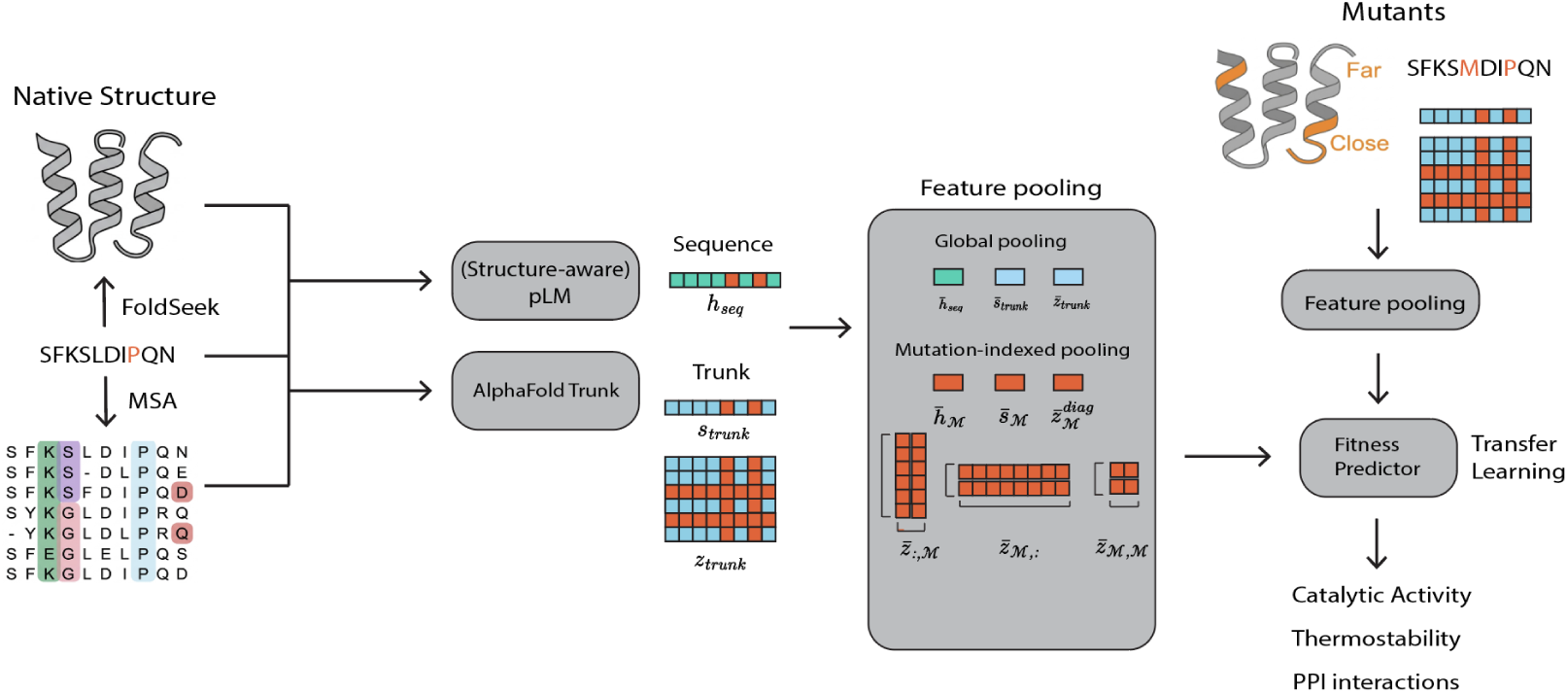
Multi-modal fitness prediction framework. A lightweight probe regresses fitness labels on pooled embeddings from two sources. Sequence embeddings come from either a sequence-only or a structure-aware pLM, the latter consuming a mutated sequence alongside the structure of its native counterpart. Trunk embeddings (added in our + *trunk* configurations), are the single and pair representations from an AlphaFold-style trunk conditioned on the sequence and its MSA. Each representation is pooled both globally and at the mutated positions, giving nine fixed-length blocks; only the three global blocks are available when the wild type of a test variant is unknown. All encoders remain frozen and only the probe is trained.

**Figure 2:**
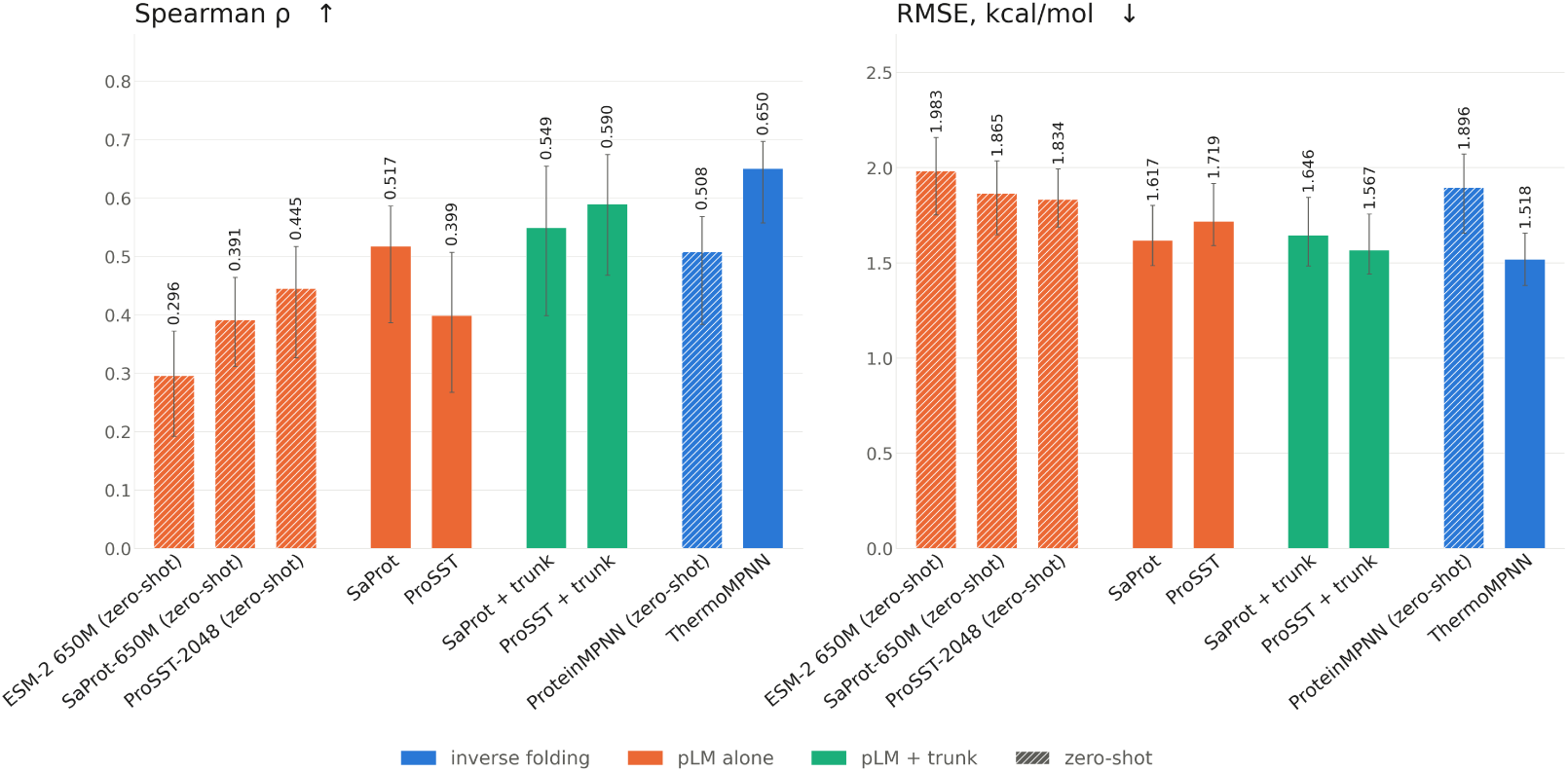
**Spearman correlation and RMSE for thermostability prediction on FireProt.**. Adding the trunk features to structure-aware pLMs such as ProSST (0.399 → 0.590) yields a larger gain than ThermoMPNN over ProteinMPNN (0.508 → 0.650). Error bars: 95% CI bootstrapped over 3 seeds.

No clear advantage emerges from folding each mutant (Table C1), so we derive structural tokens from the wild-type structure, encoded once per protein at inference.

Because the pLM and the trunk produce per-residue representations, their size grows with sequence length. To obtain a fixed-length input that can transfer to held-out sequences of any length, we pool each representation into nine blocks (Figure 1). We write *̄·* for the average over all residues (or all residue pairs) and *̄·_M_* for the average restricted to *M ⊂ {*1*, . . . , N}*, the positions at which mutants differ from their native sequences. Three out of nine blocks take the unrestricted average and are wild-type (WT) independent: *h̄*, *s̄*, and *z̄* are defined as global pooled features. The remaining six blocks restrict to *M*, which exists only relative to a corresponding wild-type sequence, and are therefore WT-dependent. Two of them pool single representations at the mutated positions, *h̄_M_* and *s̄_M_*; the other four pool the pair representation *z* over the rows, columns, diagonal entries, and submatrix indexed by *M*:

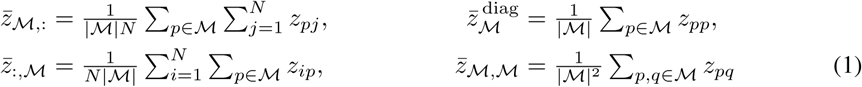

The nine pooled features are concatenated to form the covariate vector *ϕ*(*x*) *∈* R^2^*^dh^*^+2^*^ds^*^+5^*^dz^* for the fitness-prediction probe, where *d_h_* = *d*_pLM_, *d_s_* = 384, and *d_z_* = 256. For extrapolation for wild-type splits, we use only three globally pooled trunk blocks, *h̄*, *s̄*, and *z̄*. In contrast, for the other split types, we use all nine pooled block features (Figure 1). When training predictors to extrapolate to unseen backbones (wild-type splits), pLMs use only globally pooled features, i.e., *h̄*. for the other split types, both globally pooled and mutation-indexed pooled features, *h̄* and *h̄_M_*, are used.

We fit non-linear (MLP) probes on the pooled features *ϕ*(*x*) while keeping the encoder frozen, so that *ϕ* is precomputed and only the parameters of the prediction head are learned. The non-linear head is a two-layer MLP with GELU activations *σ*, formulated as *f*^^^(*x*) = *W*_2_ *σ*(*W*_1_ *ϕ*(*x*)+*b*_1_)+*b*_2_. For methods combining pretrained embeddings from pLMs and trunks, we mitigate biases from their raw magnitudes by standardizing each feature dimension using statistics based on the training split.

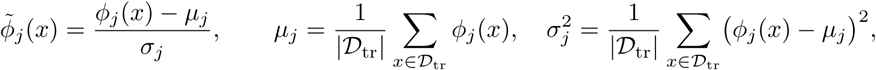

with the same *µ_j_, σ_j_* applied unchanged to the validation and test splits (see Appendix B for details on full training, and Appendix C.3 for feature standardization).

Since the trunk features pool *z* at the mutated positions, we next ask whether the trunk must be computed from the mutated sequence at all. Mutant trunks, recomputed per variant, are only slightly better on average than native trunks folded once per wild type, and the gap falls within run-to-run variation (Table C1), suggesting that most of the trunk’s contribution comes from the wild-type context rather than from variant-specific structural changes.

## 4 Experiments

**Datasets.** We evaluate on three settings: (1) FLIP2 (Didi et al., 2026), (2) the homologue-free split of FireProt (Bednar et al., 2015), designed to evaluate models trained solely on Megascale data (Tsuboyama et al., 2023), and (3) proteins with new-to-nature functions (Woolley et al., 2026). FLIP2 comprises seven datasets and 16 splits that probe generalization by withholding mutation types, mutation positions, or protein types from training. Whereas FLIP2 spans diverse protein properties and functions, FireProt isolates a single property – the thermostability of single-point mutants – and serves as a common testbed for stability predictors. We ensure that the data used to train the prediction probes do not overlap with the held-out test splits. Details of the datasets and train-test splits are provided in Appendix B.1.

Including *mutated* trunk representations in the prediction probes requires running Protenix-v2 inference for each variant, which incurs substantial computational cost. Because `TrpB` and NucB datasets contain large training sets of approximately 30k and 7k sequences, respectively, we randomly downsample `TrpB` to 10% of its original size to study the contribution of trunk representations beyond the `AMYLASE` dataset.

**Baselines and model architectures.** We compare our methods to zero-shot likelihoods, linear models, and fine-tuned pLMs, which are the baselines reported in the original FLIP2 paper (Didi et al., 2026). For our framework, we instantiate several pretrained protein encoders, including SaProt and ProSST, which consume discrete structural tokens alongside the sequence, as well as ESM-3 and multi-modal DPLM-2 (DPLM-2-mm), which consume residue tokens together with tokenized backbone structure. We also include the sequence-only variant of DPLM-2; comparing the two variants allows us to isolate the effect of the structural channel within the same architecture and pre-training recipe, so that differences can be attributed to structural input rather than to a change in model family. We additionally evaluate CHEAP embeddings (Lu et al., 2025), a compressed joint encoding of sequence and structure, to test whether a representation optimized for reconstruction retains the information required by a fitness-prediction probe. All encoders are frozen; only the prediction probe is trained.

**Metrics.** As the optimization stage in protein engineering depends on correctly ranking desired candidates rather than precise fitness prediction, we mainly report Spearman correlation. In Table C7, we additionally report normalized discounted cumulative gain (NDCG) on held-out test sets. NDCG places higher weights on high-fitness variants and thus becomes more relevant when the assays are dominated by non-functional variants (Didi et al., 2026).

### 4.1 Results on FLIP2 benchmark

For pLM-only models, we use unnormalized MLP probes, whereas pLM models with trunk features use feature-standardized MLP probes (Appendix C.3). Overall, no single configuration performs best across all datasets and splits (Table 1). Nevertheless, augmenting pLMs with structural and trunk features yields clear gains on some challenging distribution shifts. Structure-aware encoders provide their largest improvements on wild-type splits, whereas trunk features contribute most strongly on position splits. In these challenging regimes, our best configurations outperform the strongest baselines originally reported (Didi et al., 2026).

**Table 1:** Spearman correlation across FLIP2 datasets and splits, benchmarked against the baselines of Didi et al. (2026). Bold. marks the best and *^∗^* the second-best value per column. Rows without marked *↪→* refer to unnormalized MLP probes trained on only pLM, while those with marked *↪→* refer to normalized MLP probes with pLM and trunk features, and the Δ refers to gain / loss in correlation. Split names are abbreviated: *n → m* trains on *n*-point mutants and tests on *m*-point; C*→*F / F*→*C are the close-to-far and far-to-close position splits. ^a^ for the Hydro wild-type splits indicates the use of features restricted WT-free blocks (See Section 3)

| Method | Amylase |  |  |  | IREL | Hydro |  |  |  |  | Rhomax |
| --- | --- | --- | --- | --- | --- | --- | --- | --- | --- | --- | --- |
|  | Num. | Position |  | Mut. | Num. | Num. | Fit. | Wild Type<sup>a</sup> |  |  | Wild Type |
| | 1→m | c→f | f→c | by-mut | 2→m | 3→m | lo→hi | P06241 | P0A9X9 | P01053 | by-WT |
| *Sequence-input pLM* |  |  |  |  |  |  |  |  |  |  |  |
| ESM-C | 0.654 | 0.289 | 0.274 | 0.581 | **0.211** | 0.571 | 0.351 | 0.463 | 0.160 | 0.341 | 0.000 |
| $\hookrightarrow$ + trunk | 0.695 | 0.698* | 0.612* | **0.720** | 0.196 | **0.679** | 0.416* | 0.333 | 0.200 | 0.324 | 0.290 |
| $\Delta$ | +.041 | +.409 | +.338 | +.139 | -.015 | +.108 | +.065 | -.130 | +.040 | -.017 | +.290 |
| ESM2 | 0.611 | 0.284 | 0.156 | 0.594 | 0.209* | 0.520 | 0.314 | 0.424 | 0.032 | 0.129 | 0.165 |
| $\hookrightarrow$ + trunk | 0.685 | 0.665 | **0.614** | 0.701 | 0.181 | 0.659 | 0.368 | 0.275 | **0.321** | **0.413** | 0.154 |
| $\Delta$ | +.074 | +.381 | +.458 | +.107 | -.028 | +.139 | +.054 | -.149 | +.289 | +.284 | -.011 |
| DPLM-2 | 0.623 | 0.358 | 0.113 | 0.568 | 0.193 | 0.409 | 0.310 | 0.277 | 0.147 | -0.162 | 0.320 |
| $\hookrightarrow$ + trunk | 0.700 | 0.661 | 0.593 | 0.704 | 0.175 | 0.583 | 0.350 | 0.229 | 0.048 | 0.247 | 0.216 |
| $\Delta$ | +.077 | +.303 | +.480 | +.136 | -.018 | +.174 | +.040 | -.048 | -.099 | +.409 | -.104 |
| *Joint sequence-structure pLM* |  |  |  |  |  |  |  |  |  |  |  |
| CHEAP | 0.648 | 0.347 | 0.347 | 0.588 | 0.200 | 0.492 | 0.304 | 0.408 | 0.180 | 0.246 | **0.509** |
| $\hookrightarrow$ + trunk | 0.661 | 0.645 | 0.588 | 0.693 | 0.199 | 0.606 | 0.347 | 0.407 | 0.077 | 0.214 | 0.409* |
| $\Delta$ | +.013 | +.298 | +.241 | +.105 | -.001 | +.114 | +.043 | -.001 | -.103 | -.032 | -.100 |
| *Structure-input pLM* |  |  |  |  |  |  |  |  |  |  |  |
| SaProt | 0.679 | 0.527 | 0.378 | 0.636 | 0.209* | 0.577 | 0.368 | **0.496** | 0.126 | 0.202 | 0.114 |
| $\hookrightarrow$ + trunk | 0.695 | 0.670 | 0.591 | 0.690 | 0.186 | 0.662* | **0.441** | 0.269 | 0.289 | 0.393* | 0.262 |
| $\Delta$ | +.016 | +.143 | +.213 | +.054 | -.023 | +.085 | +.073 | -.227 | +.163 | +.191 | +.148 |
| ProSST | 0.632 | 0.439 | 0.339 | 0.647 | 0.183 | 0.394 | 0.268 | 0.202 | 0.063 | 0.007 | 0.025 |
| $\hookrightarrow$ + trunk | 0.678 | 0.663 | 0.602 | 0.705* | 0.171 | 0.547 | 0.332 | 0.315 | 0.227 | 0.350 | 0.031 |
| $\Delta$ | +.046 | +.224 | +.263 | +.058 | -.012 | +.153 | +.064 | +.113 | +.164 | +.343 | +.006 |
| ESM-3 | 0.689 | 0.383 | 0.398 | 0.660 | 0.137 | 0.521 | 0.348 | 0.015 | 0.166 | 0.032 | 0.192 |
| $\hookrightarrow$ + trunk | **0.733** | **0.712** | 0.605 | 0.702 | 0.179 | 0.658 | 0.397 | 0.287 | 0.311* | 0.349 | 0.315 |
| $\Delta$ | +.044 | +.329 | +.207 | +.042 | +.042 | +.137 | +.049 | +.272 | +.145 | +.317 | +.123 |
| multi-modal DPLM-2 | 0.637 | 0.423 | 0.205 | 0.574 | 0.193 | 0.392 | 0.288 | 0.419 | 0.102 | -0.043 | 0.225 |
| $\hookrightarrow$ + trunk | 0.703* | 0.647 | 0.604 | 0.703 | 0.171 | 0.616 | 0.346 | 0.480* | 0.144 | 0.190 | 0.295 |
| $\Delta$ | +.066 | +.224 | +.399 | +.129 | -.022 | +.224 | +.058 | +.061 | +.042 | +.233 | +.070 |
| Best ours | 0.733 | 0.712 | 0.614 | 0.720 | 0.211 | 0.679 | 0.441 | 0.496 | 0.321 | 0.413 | 0.509 |
| Best (Didi et al., 2026) | 0.597 | 0.617 | 0.598 | 0.681 | 0.211 | 0.604 | 0.308 | 0.426 | 0.151 | 0.329 | 0.379 |
| $\Delta$ | +.136 | +.095 | +.016 | +.039 | +.000 | +.075 | +.133 | +.070 | +.170 | +.084 | +.130 |

**Structure-aware models establish a stronger performance baseline.** Structure-aware pLMs improve Spearman correlation over sequence-only pLMs on `AMYLASE` position splits. SaProt, which uses the native backbone as input, achieves a Spearman correlation of 0.496 on the P06241 split of `HYDRO`, while CHEAP embeddings jointly trained to decode sequence and structure reach 0.509 on `RHOMAX`, suggesting that structural priors contribute to generalization across distinct protein types. As position and wild-type splits assess extrapolating protein functions governed by different structural contexts, structural information offers a clear advantage. However, the pattern reverses where structure is less informative. On `RHOMAX`, sequence-only pLMs are competitive or better, consistent with these targets containing more disordered regions and lower structural confidence (Table C2). `IRED` is a challenging case where no pLM improves over the baselines.

**Trunk features contribute to generalizing over unseen mutated positions.** On `AMYLASE`, the trunk materially contributes on position splits (close-to-far rises on average from 0.38 to 0.67 and far-to-close from 0.28 to 0.60 across pLMs), adding a modest average of +0.05 on the number split and +0.10 on the mutation split. ESM-3 with trunk is best overall, achieving the highest correlation. The same effect on the position split appears on `TrpB` subset (Table 3), where baselines struggle more and trunk features improve SaProt by 0.08, but offer no benefit to ESM-C. However, for some pLMs, such as ESM-C and SaProt on the P06241 wild-type split, the trunk can contribute negatively, suggesting that representations useful for extrapolating to unseen positions do not necessarily transfer to unseen backbones. We examine this distinction through feature and MSA ablations in Section 5.

**Table 2:** In-distribution versus out-of-distribution gains of Spearman correlation from trunk features. Gain is defined as the performance improvement from adding trunk features. Gain OOD exceeding Gain ID indicates robust extrapolation rather than better fitting

| Type | Dataset | Split | Gain ID | Gain OOD |
| --- | --- | --- | --- | --- |
| position | amylase | close-to-far | +0.073 | +0.282 |
|  |  | far-to-close | +0.050 | +0.323 |
| mutation | amylase | by-mutation | +0.076 | +0.099 |
| number | amylase | one-to-many | +0.075 | +0.063 |
|  | pdz3 | single-to-double | +0.131 | +0.042 |
|  | ired | two-to-many | +0.013 | -0.011 |
| wild-type | rhomax | by-wild-type | +0.261 | +0.075 |

**Table 3:** Trunk contribution on the TrpB subset. Spearman correlation of MLP probe. The *mutated* trunk helps the structure-aware encoder but not the sequence-only one.

| Split | Encoder | $\rho$ | $\Delta$ trunk |
| --- | --- | --- | --- |
| by-position | SaProt | 0.236 | +0.083 |
| | $\hookrightarrow$ + trunk | **0.319** | |
|  | ESM-C | **0.332** | -0.051 |
| | $\hookrightarrow$ + trunk | 0.280 | |

### 4.2 Results on FireProt benchmark

On FireProt benchmark, we compare our methods to inverse-folding methods, ThermoMPNN (Dieckhaus et al., 2024), a stability predictor built on an inverse-folding encoder of ProteinMPNN (Dauparas et al., 2022). When trained on Megascale (Tsuboyama et al., 2023) on which ThermoMPNN is fine-tuned, ProSST with trunk yields the second-best performance on both Spearman correlation and RMSE but falls behind ThermoMPNN by 0.06 correlation and 0.049 RMSE (Figure 2). The results are consistent with how zero-shot likelihood scores from ProteinMPNN are better correlated with thermostability than those from ProSST embeddings. In the transfer learning scheme, adding the trunk to ProSST yields a larger incremental gain (0.399 *→* 0.590) than ThermoMPNN improves over ProteinMPNN (0.508 *→* 0.650).

### 4.3 Extrapolation to new-to-nature functions

Woolley et al. (2026) curates proteins whose mutations confer new-to-nature functions, placing them outside the evolutionary context that pLMs have learned during training. Figure 3 shows that this regime reveals the failure of pLMs; hybrid ESM-3, though it accounts for both sequence and structure input tokens, struggles to distinguish function-enhancing mutations. However, adding trunk features to ESM-3 improves overall correlation (*−*0.60 *→ −*0.15) but does not restore a positive ranking. SaProt is better behaved, improving from (+0.17 *→* +0.35) with the trunk, although the gain is concentrated among high-fitness variants rather than distributed across the range. Trunk features therefore mitigate, but do not remove, the failure of extrapolation beyond known functions.

**Figure 3:**
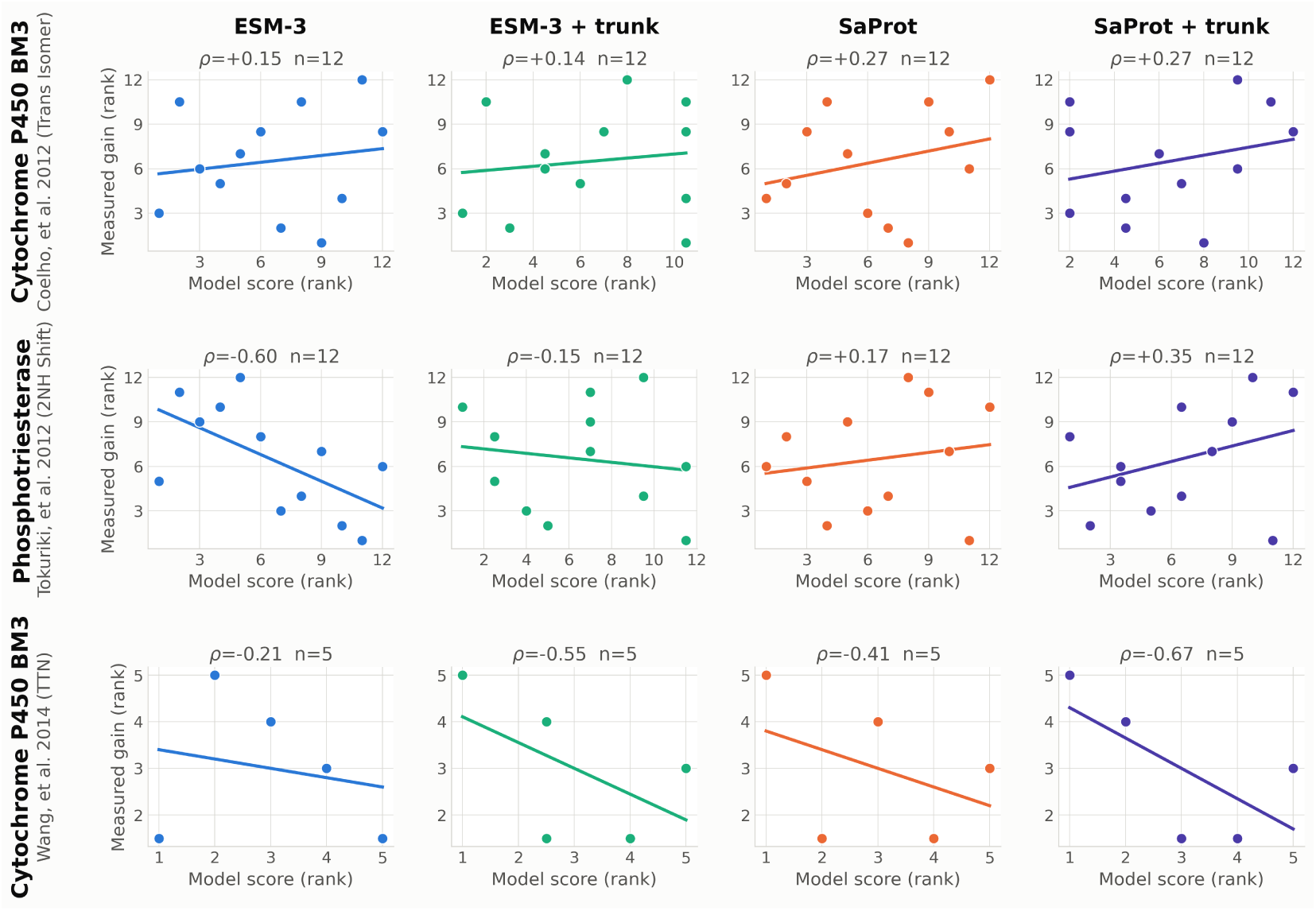
Spearman correlation between predicted variant score and measured functional gain on proteins with new-to-nature function mutations. Each row refers to each sequence model or its combination with trunk features. SaProt with trunk attains the largest positive correlation.

### 4.4 Which representations contribute to generalization?

Benchmark performance alone cannot distinguish features that are merely *informative*, correlating with fitness within the training distribution, from those that are *extrapolative*, remaining robust to the distribution shift. We therefore separate in-distribution (ID) from out-of-distribution (OOD) performance for each method. For every dataset and split, we hold out 20% of the training set as an ID test set and fit the probe on the remaining 80%, then evaluate the fitted model on the split’s designated OOD test set. When the method raises ID accuracy without narrowing the ID–OOD gap, it is considered *informative* but not *extrapolative*. The generalized method should improve OOD performance at the same level of achieved ID performance. Trunk features particularly narrow this gap on `AMYLASE` position splits, as shown in Table 2. Therefore, we attribute the trunk’s benefit to positional extrapolation: it encodes how a residue’s structural environment relates to function, which transfers to positions unseen during training but not to unseen protein backbones.

Beyond the extrapolation analysis, we ask how generalization scales with the size of the training set. We vary the number of training labels used to train the prediction head and evaluate on various FLIP2 splits. Figure C1 shows that held-out performance only slightly improves with training set size on `AMYLASE` and `HYDRO` datasets. On the `AMYLASE` position splits, adding trunk features steepens this curve, yielding a steeper scaling trend than sequence embeddings alone. On the `RHOMAX` wild-type split, which spans several backbones, standard errors are substantially wider than elsewhere, indicating that performance depends heavily on which variants are seen during training.

## 5 Ablation study

**Mutation-indexed trunk features require knowledge of the corresponding wild-type sequence.** To isolate each modality’s contribution to generalizing mutation effects, we ablate the pLM and trunk features, fitting the prediction head on each subset (Table 5). On `AMYLASE`, we quantify the contribution of mutation-indexed features by comparing performance with and without the WT-dependent blocks. For ESM-C, excluding these blocks reduces performance. On the other hand, mutation-indexed trunk features yield positive gains by +0.011 to +0.078. Therefore, the knowledge about the wild type corresponding to a certain mutant is useful when we leverage both pLMs and trunks, but not necessarily for using pre-trained embeddings from sequence-only pLMs.

**Table 4:** Co-evolutionary contribution to trunk gains on AMYLASE. Mean change in Spearman correlation over the pLM baseline, averaged across six encoders: ESM-C, ESM2, DPLM-2, SaProt, ESM-3, and multi-modal DPLM-2. MSA-PF takes the same MSA as Protenix-v2 but incorporates no geometric context. Relative recovery is Δ_MSA-PF_*/*Δ_trunk_.

| Split | $\Delta$ MSA-PF | $\Delta$ Trunk | Rel. recovery |
| --- | --- | --- | --- |
| $1 \rightarrow m$ | +0.054 | +0.016 | 3.37 |
| $c \rightarrow f$ | +0.129 | +0.201 | 0.61 |
| $f \rightarrow c$ | +0.108 | +0.159 | 0.70 |
| by-mutation | +0.022 | +0.048 | 0.43 |

**Table 5:**
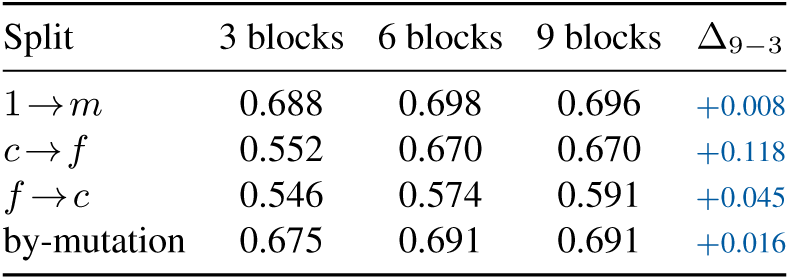
Effect of wild-type-dependent features on AMYLASE. *WT-free* blocks average over the whole chain; *WT-dependent* blocks averaged at the mutated positions. Δ is WT-full *−* WT-free, so positive values indicate that mutation-indexed pooling helps.

| Split | 3 blocks | 6 blocks | 9 blocks | $\Delta_{9-3}$ |
| --- | --- | --- | --- | --- |
| $1 \rightarrow m$ | 0.688 | 0.698 | 0.696 | +0.008 |
| $c \rightarrow f$ | 0.552 | 0.670 | 0.670 | +0.118 |
| $f \rightarrow c$ | 0.546 | 0.574 | 0.591 | +0.045 |
| by-mutation | 0.675 | 0.691 | 0.691 | +0.016 |

**Co-evolutionary signal explains a substantial fraction of the trunk gain on `amylase`**. Since the trunk encodes an MSA as input, its gain may arise partly from co-evolutionary information. To separate the contribution of co-evolutionary information from the geometric information jointly encoded in the trunk, we use MSA-Pairformer (MSA-PF) representations instead of Protenix-v2’s trunk given the same MSA. Table 4 shows that MSA-PF exceeds the trunk gain on the number split, recovers approximately 60–70% of the trunk gain on the two position splits, and approximately 43% on the mutation split. Notably, the full trunk provides the largest additional improvement on the two position splits, consistent with the geometry-aware trunk supplying information beyond co-evolutionary signals for extrapolation to unseen positions. We report the results for the other datasets and splits in Appendix D.1, which show weaker effects.

## 6 Discussion

Supervised probes trained on pLM embeddings, or even simple one-hot features, serve as strong baselines on generalization benchmarks for fitness prediction. However, their generalization performance is limited when extrapolating to new positions or entirely new protein structures. Existing studies suggest that neither scaling pLMs nor refining language architectures (Semlani et al., 2026) achieves meaningful gains, indicating a ceiling for current pLM transfer learning.

This work explores how biophysical representations, such as structural and co-evolutionary information, enable fitness predictors to overcome the bottleneck experienced by pLMs. Across the FLIP2 evaluation suite, structure-aware pLMs improve the performance baseline, particularly when extrapolating to unseen protein contexts, such as wild-type splits that benefit from structural priors. Geometric and co-evolutionary features extracted from AlphaFold-style trunks push the generalization bar, especially on position splits. The decomposition attributes this gain to greater robustness to variant-effect distribution shifts, reflecting extrapolation rather than a better fit to the training data. The benefits of combining structure-aware pLMs with trunk representations also transfer to two settings outside FLIP2: thermostability prediction on FireProt and new-to-nature protein functions.

Despite these demonstrated benefits, no single representation consistently performs best across the evaluation suite. Strong sequence-only pLMs remain more competitive in domains containing a higher proportion of intrinsically disordered proteins. Rather than proposing a one-size-fits-all method, we provide an in-depth perspective on which representations are most suitable for constructing fitness oracles for downstream tasks, given the protein families or domains of interest.

The practical value of a generalizable fitness predictor also extends to the generative design loop. ProteinDPO (Widatalla et al., 2026) exemplifies one form of this loop, adapting a pLM to the experimental characteristics of a specific campaign through Direct Preference Optimization (DPO) over binary positive and negative labels. A predictor that generalizes across mutation regimes could further relax this constraint by estimating the relative fitness of candidate sequences and providing ranked preference labels. Since our results indicate that structure-aware and trunk representations provide priors that improve generalization, a natural next step is to integrate them into post-training adaptation techniques, shifting the predictor’s role from scoring candidates under a particular fitness definition toward steering the generator across multiple design objectives, thereby better reflecting the iterative nature of real protein engineering cycles.

## 7 Limitations and Future works

Our study has several limitations that motivate future works. First, our framework relies on two pre-trained foundation models, a pLM and an AlphaFold-style trunk; extracting trunk representations is computationally expensive at inference time. As explored in Hou et al. (2026), an alternative direction is to train a single pLM to capture structural, co-evolutionary, and other biophysical information, which would reduce inference cost at the expense of greater pre-training cost. Second, our wild-type-dependent feature blocks require knowledge of the mutation positions relative to a corresponding wild-type sequence. These features are therefore unavailable under wild-type shifts where test variants originate from previously unseen wild types and the corresponding mutation set cannot be identified. Finally, predictive performance remains limited on `IRED`, where all evaluated representations achieve relatively low correlations. This suggests that the representations considered here may not fully capture necessary features for predicting IRED activity.

## Acknowledgements

**NV** and **FG** are funded by the Dutch Research Council (NWO) under the grant VI.Vidi.233.164 of the research programme Vidi ENW (https://doi.org/10.61686/IVDFS18985). **MP** is funded by a joint studentship between The University of Manchester and the A*STAR Research Attachment Programme (ARAP). This work used the Dutch national e-infrastructure with the support of the SURF Cooperative using grant no. EINF-16290.

## AI USE STATEMENT

We used generative AI tools to assist with code implementation, visualization, and the polishing or paraphrasing of written text. We did not use generative AI to develop the methods, draft the content of the paper, or cite related literature. All AI-assisted code and paraphrased text were carefully reviewed and verified by two authors. For example, LLM-generated code and visualizations were independently checked by two authors. We take full responsibility for the final content of this work, including all text, claims, code, and other artifacts produced with the assistance of generative AI tools.

## Ethics statement

We do not have a particular ethical concern about the paper, as it does not involve harmful insights, applications or human subjects. The paper has no conflicts of interest and sponsorship, discrimination/bias/fairness concerns, privacy and security issues, and legal compliance issues.

## Reproducibility statement

To facilitate the reproducibility of our research, we will release our code and results upon acceptance of the paper.

## Appendix

### A Impact statement

This work advances computational methods for protein fitness prediction, with direct applications to enzyme engineering, biocatalysis, and therapeutic protein design, where reliable ranking of candidate variants reduces the number of costly wet-lab assays required to find improved sequences.

### B Experiment details

#### B.1 Dataset details

**Fitness (FLIP2).** FLIP2 (Didi et al., 2026) extends the FLIP benchmark (Dallago et al., 2021) to span larger fitness landscapes. It has 7 datasets with 16 splits categorized into 5 extrapolation tasks. Each split holds out test variants along a different axis: (1) *Number*: train on variants with fewer mutations and test on variants with more mutations; (2) *Position*: train and test on the variants with different mutation positions; (3) *Mutation*: train and test on variants with different unique mutation types; (4) *Fitness*: train on variants with lower fitness and test on those with higher fitness; (5) *Wild Type*: train and test on variants with different sequences and structural backbones. We train the supervised probes on the same size of each dataset and split as detailed in the original Didi et al. (2026) paper.

**Amylase** Alpha amylases hydrolyse starch and are used in laundry detergents. The dataset of van der Flier et al. (2024) contains 3,706 variants of *Bacillus subtilis* alpha amylase with up to eight substitutions (488 singles). The fitness of this dataset is stain-removal activity, i.e. the enzyme performance at degrading starch-based stains under detergent conditions. This dataset includes the number split (`one-to-many`), position splits (`close-to-far` and `far-to-close`), and mutation type splits (`by-mutation`). Whether it is close or far is defined by 7.3 Å distance from active-site residues.

**Imine reductase (IRED).** IREDs reduce imines to chiral amines for pharmaceutical synthesis. Gantz et al. (2024) screened an error-prone PCR library of *Streptosporangium roseum* IRED in microfluidic droplets, yielding 17,143 variants (1,207 singles, 3,200 doubles, up to 15 mutations). Fitness is the continuous droplet-screen readout of catalytic activity on the imine substrate. This dataset includes the number split (`two-to-many`).

**Nuclease B (NucB).** NucB from *Bacillus licheniformis* degrades extracellular DNA and is of interest for disrupting biofilms in chronic wounds, but the wild type loses activity at physiological pH. Thomas et al. (2025) assayed 55,760 error-prone PCR variants at pH 7. To suppress assay noise, fitness is discretised into four ordinal activity bins: inactive (0), below wild type (1), above wild type but below the best single mutant (2), and above the best single mutant (3). This dataset includes the number split (`two-to-many`).

**Tryptophan synthase** *β***-subunit (`TrpB`).** `TrpB` synthesises tryptophan from indole and serine, an essential function that permits a growth-based selection assay; fitness is the growth enrichment of each variant, a proxy for in vivo synthesis flux. Dallago et al. (2021) measured combinatorially complete landscapes across 20 positions: nine 3-site landscapes (*∼*8,000 variants each) and one 4-site landscape (*∼*160,000 variants), 228,298 variants in total. This dataset includes the number splits (`one-to-many` and `two-to-many`) and the position split (`by-position`).

**Hydrophobic core (Hydro).** Escobedo et al. (2024) randomised seven buried core residues in each of three unrelated proteins (UniProt `P06241`, `P01053`, `P0A9X9`) to F, I, L, M, and V, assaying 24,935 variants. Fitness is thermodynamic folding stability rather than activity, reported on a common scale across scaffolds (median *−*3.21). This dataset includes the number splits (`three-to-many`), the fitness value split (`low-to-high`), and the wild-type splits (`to-P06241`, `to-P0A9X9`, `to-P01053`) that train on two scaffolds and test on the held-out third.

**Rhodopsin (Rhomax).** Microbial rhodopsins are light-activated membrane proteins used in optogenetics, where red-shifted variants are preferred. The landscape comprises 884 variants derived from 75 rhodopsin parents (Karasuyama et al., 2018; Inoue et al., 2021; Sela et al., 2024), spanning 1–6 substitutions and chimeras in which segments of other wild types replace the homologous region. Fitness is the peak absorption wavelength *λ*_max_ (nm), a spectroscopic property rather than an activity or stability measure. This dataset includes the wild-type splits (`by-wild-type`), training on 5 most common wild types and testing on 36 held-out ones.

**PDZ domain (PDZ3).** PDZ domains bind short linear motifs, often within intrinsically disordered regions. Zarin & Lehner (2024) measured binding between mutants of the PDZ3 domain and mutants of a nine-residue SLiM from CRIPT across more than 200,000 double-mutant combinations. Fitness is the binding affinity of a *pair* of sequences, making this the only two-protein landscape in FLIP2. The `single-to-double` split trains on PDZ3 single mutants and tests on 579 double mutants (one mutation in each partner) filtered for significantly non-additive binding, concentrating evaluation on epistasis.

#### B.2 Baselines

FLIP2 baselines evaluate zero-shot protein language model (pLM) sequence-likelihood scores, ridge regression supervised on pLM embeddings, and fine-tuned pLMs with a trained prediction head.

**Zero-shot sequence-likelihood scoring.** This approach requires no labelled fitness and measures the likelihood of a mutated residue relative to its corresponding wild type. For masked language models (MLM) such as CARP-640M and ESM2-650M, we can formulate a variant with mutated positions *P* as follows

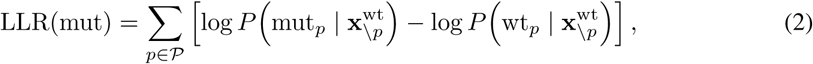

where **x**^wt^ is the wildtype sequence with position *p* masked. As an autoregressive model, Dayhoff scores are averaged over N-to-C and C-to-N decoding directions across two model variants.

**Supervised ridge regression.** Ridge regression is trained on either one-hot encoded (OHE) sequences, zero-padded to the maximum length in the dataset or the one-hot features with the zero-shot likelihood scores from pLMs. This simple supervised baseline also fits an independent additive weight to each residue-position pair, capturing the additive component of a landscape.

**Fine-tuned pLMs.** As fine-tuned pLM baselines, CARP-640M and ESMC-300M are fine-tuned on the training set of each split. For each model, the baselines consider fine-tuning from both pretrained weights and randomly initialized weights with the same architecture. For benchmarking against our method, we report the performance averaged over five random seeds for each fine-tuning configuration.

#### B.3 Hyperparameter Settings

As our methods operate in a supervised learning setting, we describe the training hyperparameters used for both linear and non-linear fitness prediction probes. For the linear probe, we use ridge regression with a fixed regularization strength of *α* = 10. For the non-linear MLP probe, we optimize the model with Adam (Kingma & Ba, 2014) using a learning rate of 10*^−^*^4^ and mean squared error (MSE) loss. We use a `LambdaLR` schedule with a linear warm-up over the first 1,000 steps, followed by a constant learning rate. The remaining hyperparameters are a batch size of 16 and 500 training epochs. Non-linear probes are used for the results reported in the main text, while linear probes are included in Appendix C.3 to provide additional context for the choice of prediction probe.

### C Supplementary results

#### C.1#Mutant structure and trunk as inputs to pLM

Table C1 assesses whether using the native structure or a structure predicted from the mutant sequence affects held-out performance, finding only marginal differences across splits. The differences are generally within *±*0.02, except SaProt on *c → f* (+0.046), where the baseline is weakest and therefore has the most room for improvement. On `amylase`, comparing native and mutated trunk features yields an even smaller difference, within *±*0.014. Overall, these results suggest that the generalizability of fitness predictors is largely invariant to whether the input structure is native or predicted from the mutant sequence. The same trend is observed when comparing fitness predictions based on *native* versus *mutant* trunk features.

**Figure C1:**
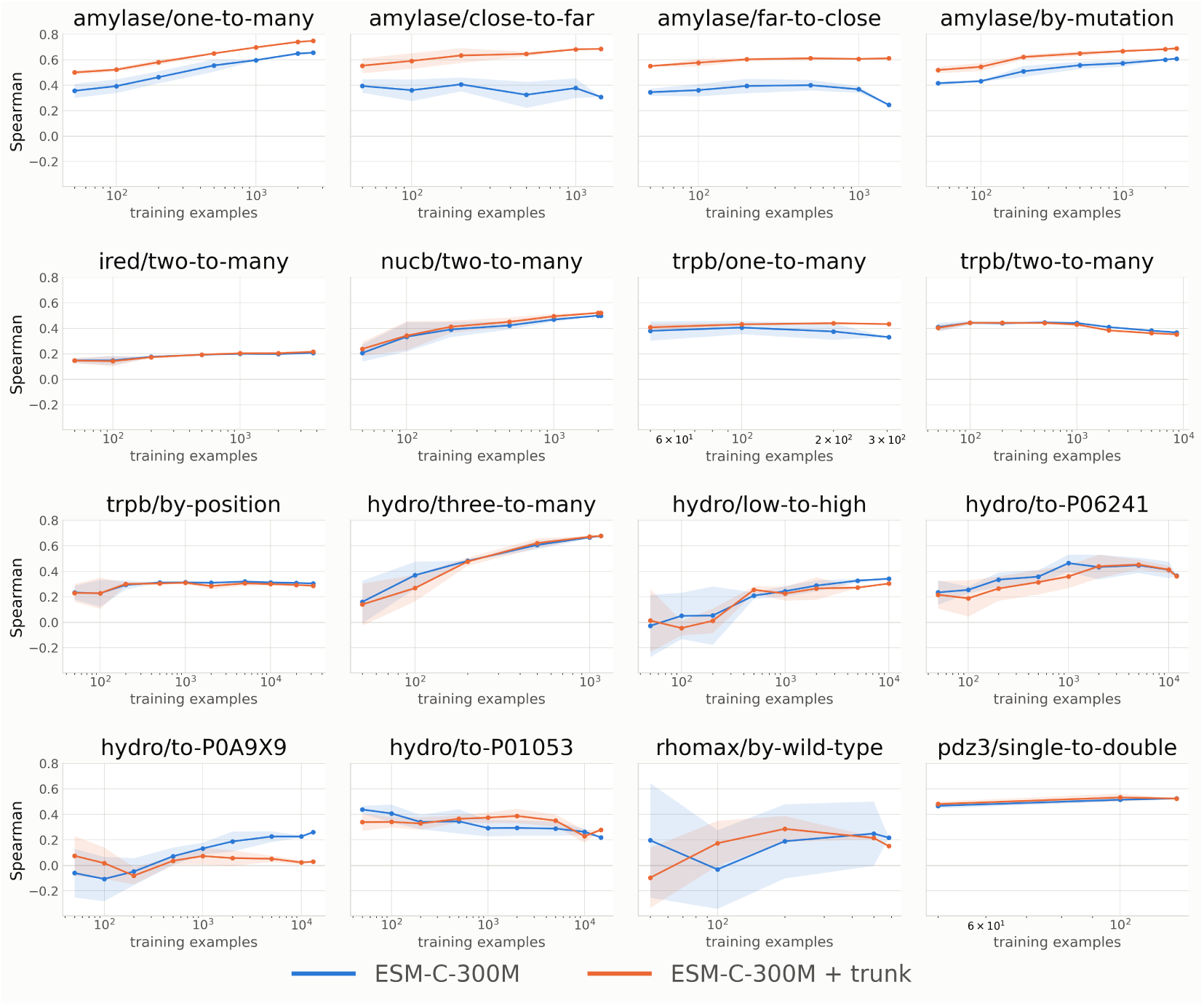
Held-out performance as a function of training set size, by FLIP2 dataset and split.

**Table C1:** Native versus mutant structure conditioning, best of ridge/MLP per cell. Δ is mutant *−* native.

| Split | Method | pLM |  |  | pLM + trunk |  |  |
| --- | --- | --- | --- | --- | --- | --- | --- |
| | | native structure | mutant structure | $\Delta$ | native trunk | mutated trunk | $\Delta$ |
| $1 \rightarrow m$ | SaProt | 0.677 | 0.679 | +0.002 | 0.747 | 0.744 | -0.003 |
|  | ProSST | 0.675 | 0.668 | -0.006 | 0.745 | 0.738 | -0.007 |
| $c \rightarrow f$ | SaProt | 0.439 | 0.485 | +0.046 | 0.677 | 0.679 | +0.002 |
|  | ProSST | 0.469 | 0.458 | -0.011 | 0.663 | 0.663 | -0.000 |
| $f \rightarrow c$ | SaProt | 0.431 | 0.445 | +0.014 | 0.593 | 0.594 | +0.001 |
|  | ProSST | 0.434 | 0.418 | -0.017 | 0.596 | 0.601 | +0.005 |
| by-mut | SaProt | 0.664 | 0.640 | -0.024 | 0.693 | 0.685 | -0.008 |
|  | ProSST | 0.681 | 0.662 | -0.019 | 0.709 | 0.695 | -0.014 |

#### C.2 Structural confidence of A M Y L A S E and R H O M A X

In Table C2, structural confidence may influence how effectively trunk representations transfer to fitness prediction. `amylase` has highly confident predicted structures (mean pLDDT 97.9; only 1.4% of residues below 90), whereas `rhomax` has substantially lower confidence (mean pLDDT 82.0; 45.2% below 90 and 30.0% below 70). This contrast suggests that the generalizability of trunk features may depend on structural quality, with less reliable structural predictions potentially limiting the transferable signal captured by the trunk.

**Table C2:** Structure quality of the two datasets. Per-residue pLDDT statistics from the predicted structures.

| Dataset | Seq. length | Mean pLDDT | % < 90 | % < 70 |
| --- | --- | --- | --- | --- |
| `amylase` | 425 | 97.9 | 1.4% | 0.5% |
| `rhomax` | 261 | 82.0 | 45.2% | 30.0% |

#### C.3 Feature Standardization and Probe Selection

The representations used in our framework come from different pretrained encoders and can therefore differ substantially in scale across feature dimensions and modalities. We study how feature standardization affects downstream fitness prediction for both ridge and MLP probes. For the linear probe, we fit ridge regression, *f*^^^(*x*) = *ϕ*(*x*)*^⊤^w*^(*α^★^*) + ^^^*b*, where the regularization strength *α^★^* is selected on the validation split according to Spearman correlation. When standardization is applied, each feature dimension is transformed as *ϕ̄_j_* (*x*) = *ϕ_j_*(*x*)*−µ_j_ /σ_j_*. Under the conventional setting, *µ_j_* and *σ_j_* are estimated from the training set and then applied unchanged to the test set. We evaluate the same standardization procedure for both ridge and MLP probes.

When combining pLM embeddings with trunk features, standardization is particularly important because the two modalities can have different raw feature scales. We therefore standardize the combined representations to a common scale to prevent differences in feature magnitude from disproportionately influencing the learned prediction weights. For pLMs without trunks, we evaluate both standardized and unstandardized features with ridge and MLP probes.

We further ask whether access to feature statistics from the test distribution can improve robustness under distribution shift. This analysis is motivated by test-time adaptation methods in computer vision, where recomputing BatchNorm statistics from batches of corrupted test samples can recover predictive performance under distribution shift (Li et al., 2018; Schneider et al., 2020; Mirza et al., 2022). Although image corruptions and protein sequence variation represent different forms of distribution shift, this motivates an analogous question in our setting: whether recomputing standardization statistics from the test distribution can improve protein fitness prediction.

Tables C3 and C4 compare standardization using training-set and test-set statistics. Recomputing the scaler from the test set does not provide a consistent improvement across datasets, encoders, or probe types. While individual configurations occasionally improve, these gains are accompanied by substantial degradations in others. Most notably, on the `HYDRO` 3 *→ m* split, test-set standardization reduces performance for every encoder under both ridge and MLP probes, with and without trunk features. These results indicate that access to test-distribution statistics alone is insufficient to reliably correct the distribution shifts considered here. We therefore use only training-set statistics whenever feature standardization is applied.

When combining pLM embeddings with trunk features, standardization is particularly important because the two modalities can have different raw feature scales. Consistent with this motivation, scaled MLP probes outperform their unscaled counterparts across the large majority of pLM+trunk configurations. For pLM-only representations, the effect of standardization is less uniform, although scaling also improves performance on average. We therefore treat standardization as particularly important when combining multiple representation modalities.

**Table C3:**
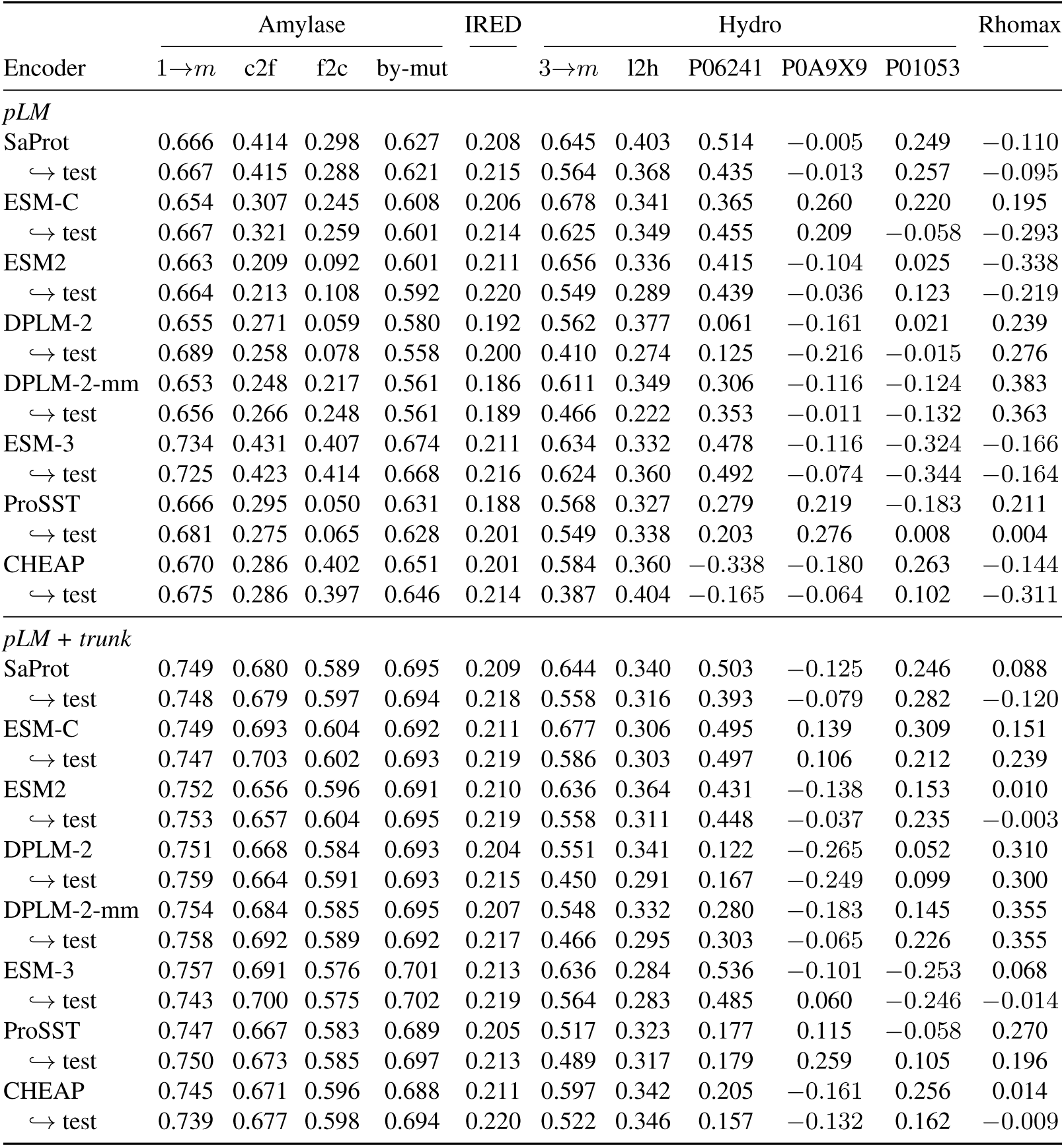
Spearman correlation of ridge probes with scaled features based on train-set vs. test-set statistics.

| Encoder | Amylase |  |  |  | IRED | Hydro |  |  |  |  | Rhomax |
| --- | --- | --- | --- | --- | --- | --- | --- | --- | --- | --- | --- |
|  | 1→m | c2f | f2c | by-mut |  | 3→m | l2h | P06241 | P0A9X9 | P01053 |  |
| *pLM* |  |  |  |  |  |  |  |  |  |  |  |
| SaProt | 0.666 | 0.414 | 0.298 | 0.627 | 0.208 | 0.645 | 0.403 | 0.514 | −0.005 | 0.249 | −0.110 |
| ↪ test | 0.667 | 0.415 | 0.288 | 0.621 | 0.215 | 0.564 | 0.368 | 0.435 | −0.013 | 0.257 | −0.095 |
| ESM-C | 0.654 | 0.307 | 0.245 | 0.608 | 0.206 | 0.678 | 0.341 | 0.365 | 0.260 | 0.220 | 0.195 |
| ↪ test | 0.667 | 0.321 | 0.259 | 0.601 | 0.214 | 0.625 | 0.349 | 0.455 | 0.209 | −0.058 | −0.293 |
| ESM2 | 0.663 | 0.209 | 0.092 | 0.601 | 0.211 | 0.656 | 0.336 | 0.415 | −0.104 | 0.025 | −0.338 |
| ↪ test | 0.664 | 0.213 | 0.108 | 0.592 | 0.220 | 0.549 | 0.289 | 0.439 | −0.036 | 0.123 | −0.219 |
| DPLM-2 | 0.655 | 0.271 | 0.059 | 0.580 | 0.192 | 0.562 | 0.377 | 0.061 | −0.161 | 0.021 | 0.239 |
| ↪ test | 0.689 | 0.258 | 0.078 | 0.558 | 0.200 | 0.410 | 0.274 | 0.125 | −0.216 | −0.015 | 0.276 |
| DPLM-2-mm | 0.653 | 0.248 | 0.217 | 0.561 | 0.186 | 0.611 | 0.349 | 0.306 | −0.116 | −0.124 | 0.383 |
| ↪ test | 0.656 | 0.266 | 0.248 | 0.561 | 0.189 | 0.466 | 0.222 | 0.353 | −0.011 | −0.132 | 0.363 |
| ESM-3 | 0.734 | 0.431 | 0.407 | 0.674 | 0.211 | 0.634 | 0.332 | 0.478 | −0.116 | −0.324 | −0.166 |
| ↪ test | 0.725 | 0.423 | 0.414 | 0.668 | 0.216 | 0.624 | 0.360 | 0.492 | −0.074 | −0.344 | −0.164 |
| ProSST | 0.666 | 0.295 | 0.050 | 0.631 | 0.188 | 0.568 | 0.327 | 0.279 | 0.219 | −0.183 | 0.211 |
| ↪ test | 0.681 | 0.275 | 0.065 | 0.628 | 0.201 | 0.549 | 0.338 | 0.203 | 0.276 | 0.008 | 0.004 |
| CHEAP | 0.670 | 0.286 | 0.402 | 0.651 | 0.201 | 0.584 | 0.360 | −0.338 | −0.180 | 0.263 | −0.144 |
| ↪ test | 0.675 | 0.286 | 0.397 | 0.646 | 0.214 | 0.387 | 0.404 | −0.165 | −0.064 | 0.102 | −0.311 |
| *pLM + trunk* |  |  |  |  |  |  |  |  |  |  |  |
| SaProt | 0.749 | 0.680 | 0.589 | 0.695 | 0.209 | 0.644 | 0.340 | 0.503 | −0.125 | 0.246 | 0.088 |
| ↪ test | 0.748 | 0.679 | 0.597 | 0.694 | 0.218 | 0.558 | 0.316 | 0.393 | −0.079 | 0.282 | −0.120 |
| ESM-C | 0.749 | 0.693 | 0.604 | 0.692 | 0.211 | 0.677 | 0.306 | 0.495 | 0.139 | 0.309 | 0.151 |
| ↪ test | 0.747 | 0.703 | 0.602 | 0.693 | 0.219 | 0.586 | 0.303 | 0.497 | 0.106 | 0.212 | 0.239 |
| ESM2 | 0.752 | 0.656 | 0.596 | 0.691 | 0.210 | 0.636 | 0.364 | 0.431 | −0.138 | 0.153 | 0.010 |
| ↪ test | 0.753 | 0.657 | 0.604 | 0.695 | 0.219 | 0.558 | 0.311 | 0.448 | −0.037 | 0.235 | −0.003 |
| DPLM-2 | 0.751 | 0.668 | 0.584 | 0.693 | 0.204 | 0.551 | 0.341 | 0.122 | −0.265 | 0.052 | 0.310 |
| ↪ test | 0.759 | 0.664 | 0.591 | 0.693 | 0.215 | 0.450 | 0.291 | 0.167 | −0.249 | 0.099 | 0.300 |
| DPLM-2-mm | 0.754 | 0.684 | 0.585 | 0.695 | 0.207 | 0.548 | 0.332 | 0.280 | −0.183 | 0.145 | 0.355 |
| ↪ test | 0.758 | 0.692 | 0.589 | 0.692 | 0.217 | 0.466 | 0.295 | 0.303 | −0.065 | 0.226 | 0.355 |
| ESM-3 | 0.757 | 0.691 | 0.576 | 0.701 | 0.213 | 0.636 | 0.284 | 0.536 | −0.101 | −0.253 | 0.068 |
| ↪ test | 0.743 | 0.700 | 0.575 | 0.702 | 0.219 | 0.564 | 0.283 | 0.485 | 0.060 | −0.246 | −0.014 |
| ProSST | 0.747 | 0.667 | 0.583 | 0.689 | 0.205 | 0.517 | 0.323 | 0.177 | 0.115 | −0.058 | 0.270 |
| ↪ test | 0.750 | 0.673 | 0.585 | 0.697 | 0.213 | 0.489 | 0.317 | 0.179 | 0.259 | 0.105 | 0.196 |
| CHEAP | 0.745 | 0.671 | 0.596 | 0.688 | 0.211 | 0.597 | 0.342 | 0.205 | −0.161 | 0.256 | 0.014 |
| ↪ test | 0.739 | 0.677 | 0.598 | 0.694 | 0.220 | 0.522 | 0.346 | 0.157 | −0.132 | 0.162 | −0.009 |

### D Ablation study

#### D.1 Disentangling Co-evolutionary and Geometric Signals

Protenix-v2 takes an MSA as input and learns representations that jointly encode co-evolutionary and geometric information. To estimate how much of the trunk benefit can be attributed to MSA-derived signal alone, we compare against an MSA Pairformer (MSA-PF) trained with masked language modelling on protocol-matched ColabFold alignments (`uniref30 2103` + `colabfold envdb 202108`), but without an explicit structural objective. MSA-PF therefore serves as a control for co-evolutionary information rather than a strict causal decomposition of the trunk representation.

For this analysis, the pLM baseline and pLM+MSA-PF are evaluated under the same model-selection procedure, choosing the better of ridge regression and MLP on the validation set. The pLM+trunk values are taken from Table 1. To ensure that the two augmentations are compared against the same reference, both Δ_MSA-PF_ and Δ_trunk_ in Table D1 are computed relative to the pLM baseline reported in that table, rather than the pLM-only values in Table 1.

**Table C4:**
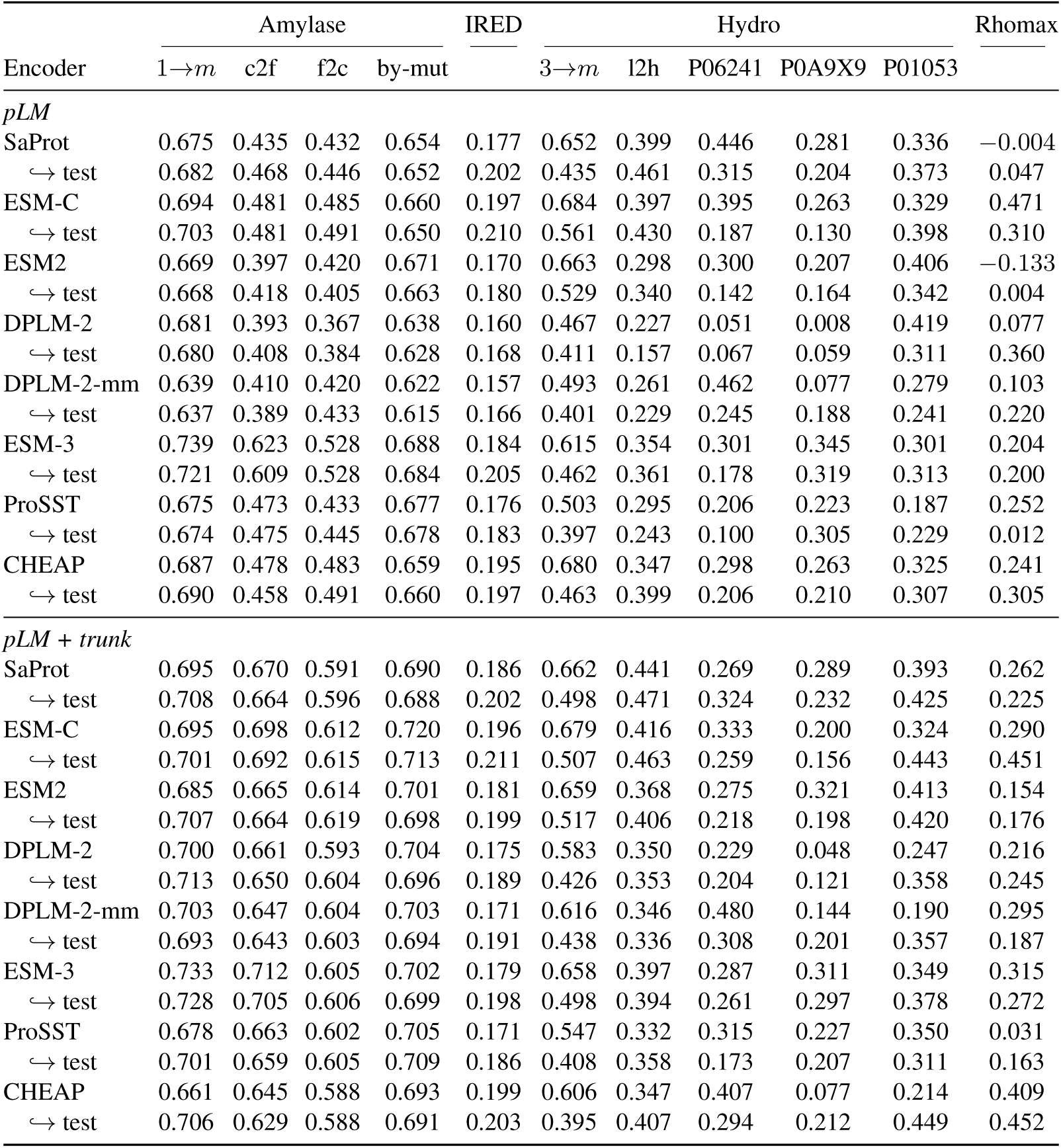
Spearman correlation of MLP probes with scaled features based on on train-set vs. test-set statistics.

|  | Amylase |  |  |  | IRED | Hydro |  |  |  |  | Rhomax |
| --- | --- | --- | --- | --- | --- | --- | --- | --- | --- | --- | --- |
| Encoder | 1→m | c2f | f2c | by-mut |  | 3→m | l2h | P06241 | P0A9X9 | P01053 |  |
| *pLM* |  |  |  |  |  |  |  |  |  |  |  |
| SaProt | 0.675 | 0.435 | 0.432 | 0.654 | 0.177 | 0.652 | 0.399 | 0.446 | 0.281 | 0.336 | −0.004 |
| ↪ test | 0.682 | 0.468 | 0.446 | 0.652 | 0.202 | 0.435 | 0.461 | 0.315 | 0.204 | 0.373 | 0.047 |
| ESM-C | 0.694 | 0.481 | 0.485 | 0.660 | 0.197 | 0.684 | 0.397 | 0.395 | 0.263 | 0.329 | 0.471 |
| ↪ test | 0.703 | 0.481 | 0.491 | 0.650 | 0.210 | 0.561 | 0.430 | 0.187 | 0.130 | 0.398 | 0.310 |
| ESM2 | 0.669 | 0.397 | 0.420 | 0.671 | 0.170 | 0.663 | 0.298 | 0.300 | 0.207 | 0.406 | −0.133 |
| ↪ test | 0.668 | 0.418 | 0.405 | 0.663 | 0.180 | 0.529 | 0.340 | 0.142 | 0.164 | 0.342 | 0.004 |
| DPLM-2 | 0.681 | 0.393 | 0.367 | 0.638 | 0.160 | 0.467 | 0.227 | 0.051 | 0.008 | 0.419 | 0.077 |
| ↪ test | 0.680 | 0.408 | 0.384 | 0.628 | 0.168 | 0.411 | 0.157 | 0.067 | 0.059 | 0.311 | 0.360 |
| DPLM-2-mm | 0.639 | 0.410 | 0.420 | 0.622 | 0.157 | 0.493 | 0.261 | 0.462 | 0.077 | 0.279 | 0.103 |
| ↪ test | 0.637 | 0.389 | 0.433 | 0.615 | 0.166 | 0.401 | 0.229 | 0.245 | 0.188 | 0.241 | 0.220 |
| ESM-3 | 0.739 | 0.623 | 0.528 | 0.688 | 0.184 | 0.615 | 0.354 | 0.301 | 0.345 | 0.301 | 0.204 |
| ↪ test | 0.721 | 0.609 | 0.528 | 0.684 | 0.205 | 0.462 | 0.361 | 0.178 | 0.319 | 0.313 | 0.200 |
| ProSST | 0.675 | 0.473 | 0.433 | 0.677 | 0.176 | 0.503 | 0.295 | 0.206 | 0.223 | 0.187 | 0.252 |
| ↪ test | 0.674 | 0.475 | 0.445 | 0.678 | 0.183 | 0.397 | 0.243 | 0.100 | 0.305 | 0.229 | 0.012 |
| CHEAP | 0.687 | 0.478 | 0.483 | 0.659 | 0.195 | 0.680 | 0.347 | 0.298 | 0.263 | 0.325 | 0.241 |
| ↪ test | 0.690 | 0.458 | 0.491 | 0.660 | 0.197 | 0.463 | 0.399 | 0.206 | 0.210 | 0.307 | 0.305 |
| *pLM + trunk* |  |  |  |  |  |  |  |  |  |  |  |
| SaProt | 0.695 | 0.670 | 0.591 | 0.690 | 0.186 | 0.662 | 0.441 | 0.269 | 0.289 | 0.393 | 0.262 |
| ↪ test | 0.708 | 0.664 | 0.596 | 0.688 | 0.202 | 0.498 | 0.471 | 0.324 | 0.232 | 0.425 | 0.225 |
| ESM-C | 0.695 | 0.698 | 0.612 | 0.720 | 0.196 | 0.679 | 0.416 | 0.333 | 0.200 | 0.324 | 0.290 |
| ↪ test | 0.701 | 0.692 | 0.615 | 0.713 | 0.211 | 0.507 | 0.463 | 0.259 | 0.156 | 0.443 | 0.451 |
| ESM2 | 0.685 | 0.665 | 0.614 | 0.701 | 0.181 | 0.659 | 0.368 | 0.275 | 0.321 | 0.413 | 0.154 |
| ↪ test | 0.707 | 0.664 | 0.619 | 0.698 | 0.199 | 0.517 | 0.406 | 0.218 | 0.198 | 0.420 | 0.176 |
| DPLM-2 | 0.700 | 0.661 | 0.593 | 0.704 | 0.175 | 0.583 | 0.350 | 0.229 | 0.048 | 0.247 | 0.216 |
| ↪ test | 0.713 | 0.650 | 0.604 | 0.696 | 0.189 | 0.426 | 0.353 | 0.204 | 0.121 | 0.358 | 0.245 |
| DPLM-2-mm | 0.703 | 0.647 | 0.604 | 0.703 | 0.171 | 0.616 | 0.346 | 0.480 | 0.144 | 0.190 | 0.295 |
| ↪ test | 0.693 | 0.643 | 0.603 | 0.694 | 0.191 | 0.438 | 0.336 | 0.308 | 0.201 | 0.357 | 0.187 |
| ESM-3 | 0.733 | 0.712 | 0.605 | 0.702 | 0.179 | 0.658 | 0.397 | 0.287 | 0.311 | 0.349 | 0.315 |
| ↪ test | 0.728 | 0.705 | 0.606 | 0.699 | 0.198 | 0.498 | 0.394 | 0.261 | 0.297 | 0.378 | 0.272 |
| ProSST | 0.678 | 0.663 | 0.602 | 0.705 | 0.171 | 0.547 | 0.332 | 0.315 | 0.227 | 0.350 | 0.031 |
| ↪ test | 0.701 | 0.659 | 0.605 | 0.709 | 0.186 | 0.408 | 0.358 | 0.173 | 0.207 | 0.311 | 0.163 |
| CHEAP | 0.661 | 0.645 | 0.588 | 0.693 | 0.199 | 0.606 | 0.347 | 0.407 | 0.077 | 0.214 | 0.409 |
| ↪ test | 0.706 | 0.629 | 0.588 | 0.691 | 0.203 | 0.395 | 0.407 | 0.294 | 0.212 | 0.449 | 0.452 |

AMYLASE provides the cleanest comparison, with six pLM encoders evaluated across all four splits. On IRED, differences among the pLM, MSA-PF, and trunk configurations remain small, within approximately 0.04 in Spearman’s *ρ*, consistent with the low performance ceiling of approximately 0.22 on this dataset. The corresponding results on HYDRO show weaker and less consistent effects than on AMYLASE.

#### D.2 Best Published FLIP2 Baselines

Table 1 reports only the best published Spearman correlation from Didi et al. (2026) for each evaluated split to keep the main benchmark compact. Table D2 lists the corresponding baseline method that achieves each value.

**Table C5:**
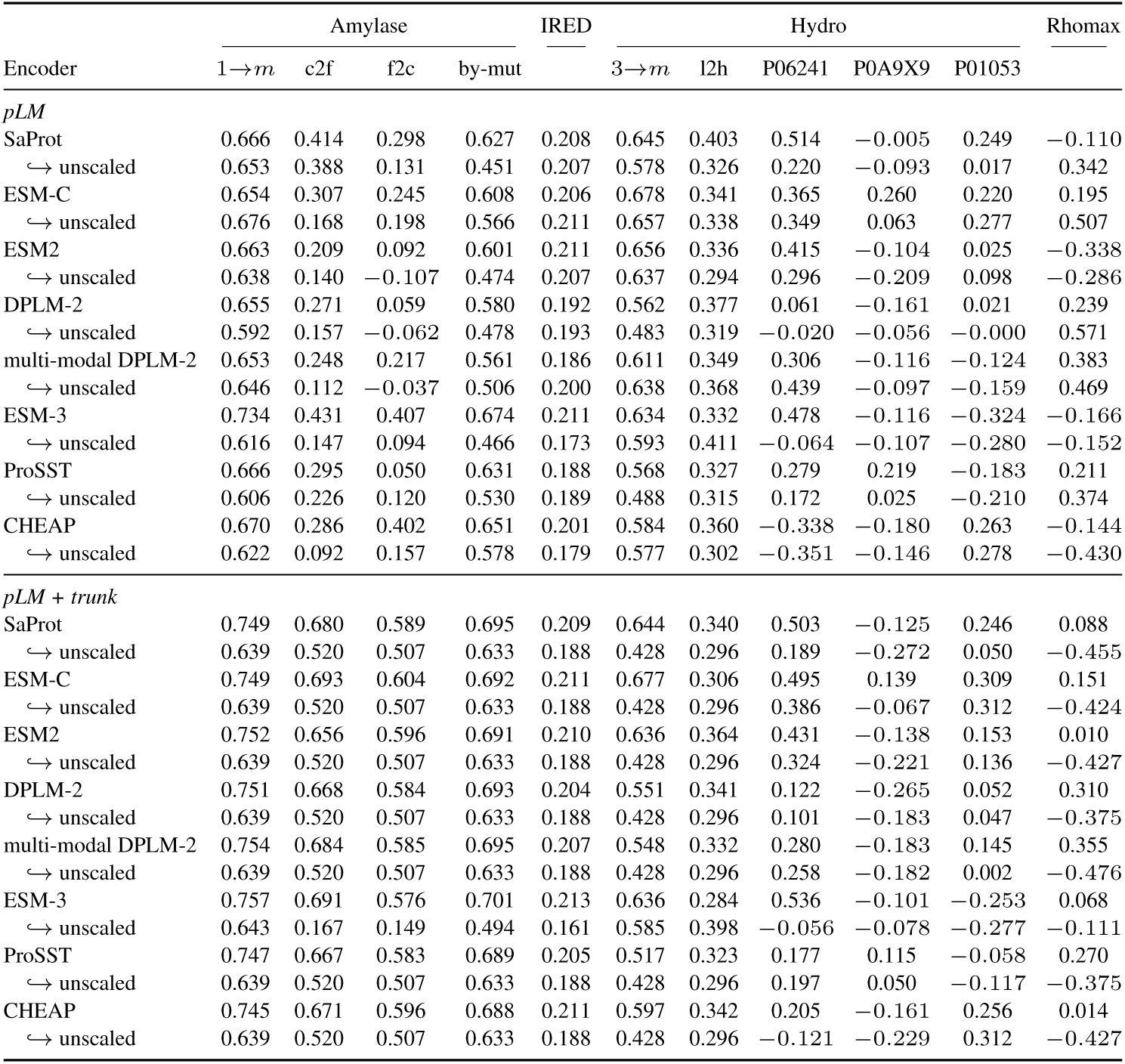
Spearman correlation of Ridge probes with scaled or unscaled features. Scaled features are standardized feature-wise using the training-set mean and standard deviation.

**Table C6:** Spearman correlation of MLP probes with scaled or unscaled features. Scaled features are standardized by their embedding dimensions.

| Encoder | Amylase |  |  |  | IRED | Hydro |  |  |  |  | Rhmax |
| --- | --- | --- | --- | --- | --- | --- | --- | --- | --- | --- | --- |
|  | 1→m | c2f | f2c | by-mut |  | 3→m | l2h | P06241 | P0A9X9 | P01053 |  |
| *pLM* |  |  |  |  |  |  |  |  |  |  |  |
| SaProt | 0.675 | 0.435 | 0.432 | 0.654 | 0.177 | 0.652 | 0.399 | 0.446 | 0.281 | 0.336 | −0.004 |
| ↪ unscaled | 0.679 | 0.527 | 0.378 | 0.636 | 0.209 | 0.577 | 0.368 | 0.496 | 0.126 | 0.202 | 0.114 |
| ESM-C | 0.694 | 0.481 | 0.485 | 0.660 | 0.197 | 0.684 | 0.397 | 0.395 | 0.263 | 0.329 | 0.471 |
| ↪ unscaled | 0.654 | 0.289 | 0.274 | 0.581 | 0.211 | 0.571 | 0.351 | 0.463 | 0.160 | 0.341 | 0.000 |
| ESM2 | 0.669 | 0.397 | 0.420 | 0.671 | 0.170 | 0.663 | 0.298 | 0.300 | 0.207 | 0.406 | −0.133 |
| ↪ unscaled | 0.611 | 0.284 | 0.156 | 0.594 | 0.209 | 0.520 | 0.314 | 0.424 | 0.032 | 0.129 | 0.165 |
| DPLM-2 | 0.681 | 0.393 | 0.367 | 0.638 | 0.160 | 0.467 | 0.227 | 0.051 | 0.008 | 0.419 | 0.077 |
| ↪ unscaled | 0.623 | 0.358 | 0.113 | 0.568 | 0.193 | 0.409 | 0.310 | 0.277 | 0.147 | −0.162 | 0.320 |
| multi-modal DPLM-2 | 0.639 | 0.410 | 0.420 | 0.622 | 0.157 | 0.493 | 0.261 | 0.462 | 0.077 | 0.279 | 0.103 |
| ↪ unscaled | 0.637 | 0.423 | 0.205 | 0.574 | 0.193 | 0.392 | 0.288 | 0.419 | 0.102 | −0.043 | 0.225 |
| ESM-3 | 0.739 | 0.623 | 0.528 | 0.688 | 0.184 | 0.615 | 0.354 | 0.301 | 0.345 | 0.301 | 0.204 |
| ↪ unscaled | 0.689 | 0.383 | 0.398 | 0.660 | 0.137 | 0.521 | 0.348 | 0.015 | 0.166 | 0.032 | 0.192 |
| ProSST | 0.675 | 0.473 | 0.433 | 0.677 | 0.176 | 0.503 | 0.295 | 0.206 | 0.223 | 0.187 | 0.252 |
| ↪ unscaled | 0.632 | 0.439 | 0.339 | 0.647 | 0.183 | 0.394 | 0.268 | 0.202 | 0.063 | 0.007 | 0.025 |
| CHEAP | 0.687 | 0.478 | 0.483 | 0.659 | 0.195 | 0.680 | 0.347 | 0.298 | 0.263 | 0.325 | 0.241 |
| ↪ unscaled | 0.648 | 0.347 | 0.347 | 0.588 | 0.200 | 0.492 | 0.304 | 0.408 | 0.180 | 0.246 | 0.509 |
| *pLM + trunk* |  |  |  |  |  |  |  |  |  |  |  |
| SaProt | 0.695 | 0.670 | 0.591 | 0.690 | 0.186 | 0.662 | 0.441 | 0.269 | 0.289 | 0.393 | 0.262 |
| ↪ unscaled | 0.636 | 0.570 | 0.527 | 0.664 | 0.185 | 0.446 | 0.179 | 0.330 | −0.243 | 0.297 | 0.148 |
| ESM-C | 0.695 | 0.698 | 0.612 | 0.720 | 0.196 | 0.679 | 0.416 | 0.333 | 0.200 | 0.324 | 0.290 |
| ↪ unscaled | 0.627 | 0.571 | 0.525 | 0.652 | 0.170 | 0.480 | 0.186 | 0.319 | −0.136 | 0.177 | 0.211 |
| ESM2 | 0.685 | 0.665 | 0.614 | 0.701 | 0.181 | 0.659 | 0.368 | 0.275 | 0.321 | 0.413 | 0.154 |
| ↪ unscaled | 0.652 | 0.565 | 0.549 | 0.667 | 0.188 | 0.456 | 0.201 | 0.334 | −0.062 | 0.377 | 0.152 |
| DPLM-2 | 0.700 | 0.661 | 0.593 | 0.704 | 0.175 | 0.583 | 0.350 | 0.229 | 0.048 | 0.247 | 0.216 |
| ↪ unscaled | 0.631 | 0.545 | 0.551 | 0.677 | 0.188 | 0.456 | 0.168 | 0.263 | −0.172 | 0.329 | 0.151 |
| multi-modal DPLM-2 | 0.703 | 0.647 | 0.604 | 0.703 | 0.171 | 0.616 | 0.346 | 0.480 | 0.144 | 0.190 | 0.295 |
| ↪ unscaled | 0.636 | 0.562 | 0.566 | 0.668 | 0.181 | 0.441 | 0.165 | 0.334 | −0.103 | 0.380 | 0.151 |
| ESM-3 | 0.733 | 0.712 | 0.605 | 0.702 | 0.179 | 0.658 | 0.397 | 0.287 | 0.311 | 0.349 | 0.315 |
| ↪ unscaled | 0.706 | 0.439 | 0.433 | 0.669 | 0.135 | 0.520 | 0.293 | 0.052 | 0.078 | 0.112 | 0.219 |
| ProSST | 0.678 | 0.663 | 0.602 | 0.705 | 0.171 | 0.547 | 0.332 | 0.315 | 0.227 | 0.350 | 0.031 |
| ↪ unscaled | 0.641 | 0.558 | 0.535 | 0.658 | 0.168 | 0.469 | 0.150 | 0.287 | −0.204 | 0.292 | −0.037 |
| CHEAP | 0.661 | 0.645 | 0.588 | 0.693 | 0.199 | 0.606 | 0.347 | 0.407 | 0.077 | 0.214 | 0.409 |
| ↪ unscaled | 0.615 | 0.543 | 0.512 | 0.653 | 0.148 | 0.468 | 0.178 | 0.326 | −0.161 | 0.322 | −0.051 |

**Table C7:**
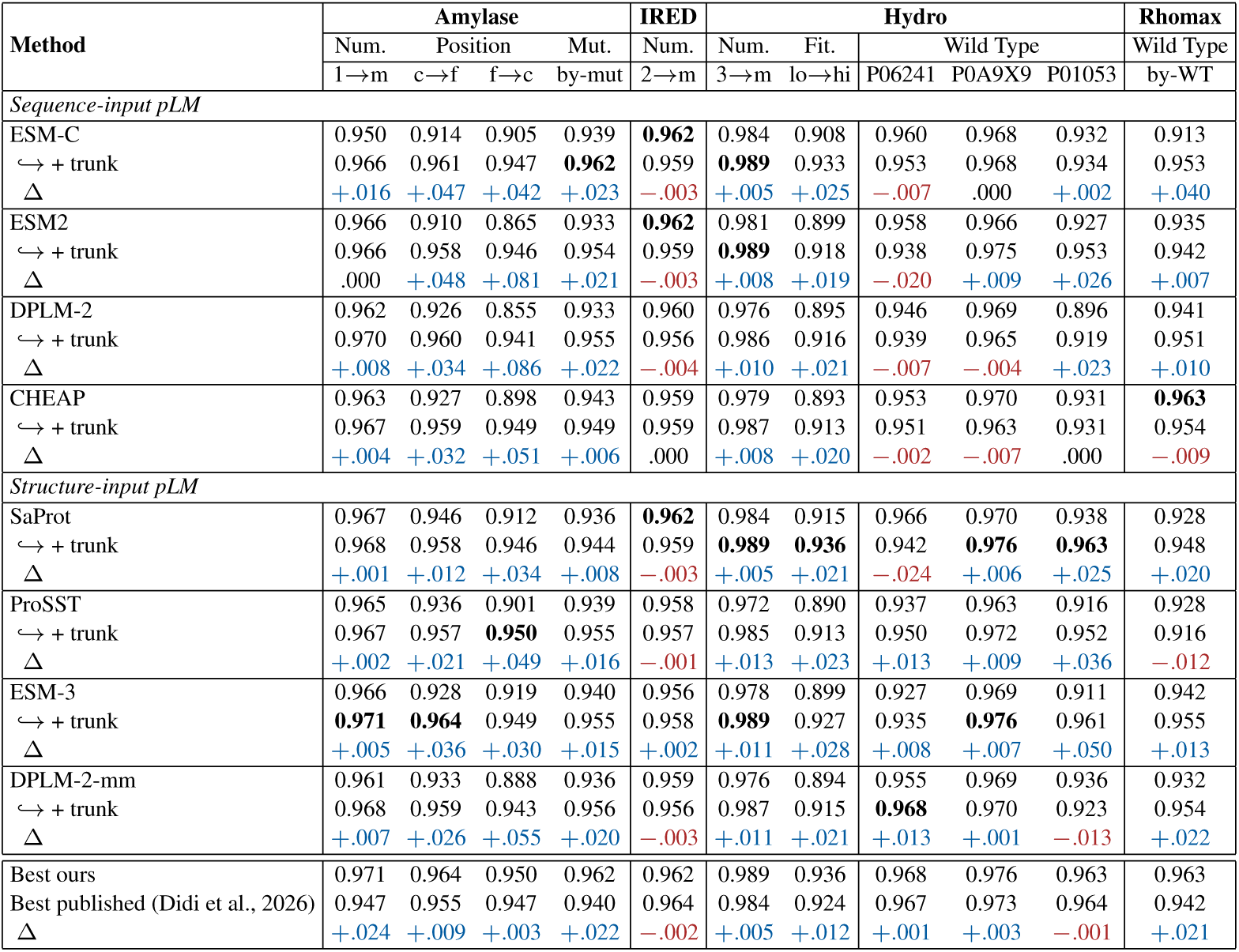
NDCG across FLIP2 datasets and splits, benchmarked against the baselines of Didi et al. (2026). Bold. marks the best value per column; where several entries tie at three decimals, all are bolded. Rows marked *↪→* augment the above encoder with Protenix-v2 trunk features, and the Δ refers to gain / loss in NDCG. Split names are abbreviated: *n→m* trains on *n*-point mutants and tests on *m*-point; C*→*F / F*→*C are the close-to-far and far-to-close position splits. Wild-type splits by protein sequences. Because NDCG is top-heavy and bounded above, values cluster near 1 and the margins are correspondingly compressed relative to Table 1. The final rows contrast the best of our models on each split with the best published value over all four baseline settings of Didi et al. (2026).

| Method | Amylase |  |  |  | IREL | Hydro |  |  |  |  | Rhomax |
| --- | --- | --- | --- | --- | --- | --- | --- | --- | --- | --- | --- |
|  | Num. | Position |  | Mut. | Num. | Num. | Fit. | Wild Type |  |  | Wild Type |
| | 1→m | c→f | f→c | by-mut | 2→m | 3→m | lo→hi | P06241 | P0A9X9 | P01053 | by-WT |
| *Sequence-input pLM* |  |  |  |  |  |  |  |  |  |  |  |
| ESM-C | 0.950 | 0.914 | 0.905 | 0.939 | **0.962** | 0.984 | 0.908 | 0.960 | 0.968 | 0.932 | 0.913 |
| $\hookrightarrow$ + trunk | 0.966 | 0.961 | 0.947 | **0.962** | 0.959 | **0.989** | 0.933 | 0.953 | 0.968 | 0.934 | 0.953 |
| $\Delta$ | **+0.016** | **+0.047** | **+0.042** | **+0.023** | **-.003** | **+0.005** | **+0.025** | **-.007** | .000 | **+0.002** | **+0.040** |
| ESM2 | 0.966 | 0.910 | 0.865 | 0.933 | **0.962** | 0.981 | 0.899 | 0.958 | 0.966 | 0.927 | 0.935 |
| $\hookrightarrow$ + trunk | 0.966 | 0.958 | 0.946 | 0.954 | 0.959 | **0.989** | 0.918 | 0.938 | 0.975 | 0.953 | 0.942 |
| $\Delta$ | .000 | **+0.048** | **+0.081** | **+0.021** | **-.003** | **+0.008** | **+0.019** | **-.020** | **+0.009** | **+0.026** | **+0.007** |
| DPLM-2 | 0.962 | 0.926 | 0.855 | 0.933 | 0.960 | 0.976 | 0.895 | 0.946 | 0.969 | 0.896 | 0.941 |
| $\hookrightarrow$ + trunk | 0.970 | 0.960 | 0.941 | 0.955 | 0.956 | 0.986 | 0.916 | 0.939 | 0.965 | 0.919 | 0.951 |
| $\Delta$ | **+0.008** | **+0.034** | **+0.086** | **+0.022** | **-.004** | **+0.010** | **+0.021** | **-.007** | **-.004** | **+0.023** | **+0.010** |
| CHEAP | 0.963 | 0.927 | 0.898 | 0.943 | 0.959 | 0.979 | 0.893 | 0.953 | 0.970 | 0.931 | **0.963** |
| $\hookrightarrow$ + trunk | 0.967 | 0.959 | 0.949 | 0.949 | 0.959 | 0.987 | 0.913 | 0.951 | 0.963 | 0.931 | 0.954 |
| $\Delta$ | **+0.004** | **+0.032** | **+0.051** | **+0.006** | .000 | **+0.008** | **+0.020** | **-.002** | **-.007** | .000 | **-.009** |
| *Structure-input pLM* |  |  |  |  |  |  |  |  |  |  |  |
| SaProt | 0.967 | 0.946 | 0.912 | 0.936 | **0.962** | 0.984 | 0.915 | 0.966 | 0.970 | 0.938 | 0.928 |
| $\hookrightarrow$ + trunk | 0.968 | 0.958 | 0.946 | 0.944 | 0.959 | **0.989** | **0.936** | 0.942 | **0.976** | **0.963** | 0.948 |
| $\Delta$ | **+0.001** | **+0.012** | **+0.034** | **+0.008** | **-.003** | **+0.005** | **+0.021** | **-.024** | **+0.006** | **+0.025** | **+0.020** |
| ProSST | 0.965 | 0.936 | 0.901 | 0.939 | 0.958 | 0.972 | 0.890 | 0.937 | 0.963 | 0.916 | 0.928 |
| $\hookrightarrow$ + trunk | 0.967 | 0.957 | **0.950** | 0.955 | 0.957 | 0.985 | 0.913 | 0.950 | 0.972 | 0.952 | 0.916 |
| $\Delta$ | **+0.002** | **+0.021** | **+0.049** | **+0.016** | **-.001** | **+0.013** | **+0.023** | **+0.013** | **+0.009** | **+0.036** | **-.012** |
| ESM-3 | 0.966 | 0.928 | 0.919 | 0.940 | 0.956 | 0.978 | 0.899 | 0.927 | 0.969 | 0.911 | 0.942 |
| $\hookrightarrow$ + trunk | **0.971** | **0.964** | 0.949 | 0.955 | 0.958 | **0.989** | 0.927 | 0.935 | **0.976** | 0.961 | 0.955 |
| $\Delta$ | **+0.005** | **+0.036** | **+0.030** | **+0.015** | **+0.002** | **+0.011** | **+0.028** | **+0.008** | **+0.007** | **+0.050** | **+0.013** |
| DPLM-2-mm | 0.961 | 0.933 | 0.888 | 0.936 | 0.959 | 0.976 | 0.894 | 0.955 | 0.969 | 0.936 | 0.932 |
| $\hookrightarrow$ + trunk | 0.968 | 0.959 | 0.943 | 0.956 | 0.956 | 0.987 | 0.915 | **0.968** | 0.970 | 0.923 | 0.954 |
| $\Delta$ | **+0.007** | **+0.026** | **+0.055** | **+0.020** | **-.003** | **+0.011** | **+0.021** | **+0.013** | **+0.001** | **-.013** | **+0.022** |
| Best ours | 0.971 | 0.964 | 0.950 | 0.962 | 0.962 | 0.989 | 0.936 | 0.968 | 0.976 | 0.963 | 0.963 |
| Best published (Didi et al., 2026) | 0.947 | 0.955 | 0.947 | 0.940 | 0.964 | 0.984 | 0.924 | 0.967 | 0.973 | 0.964 | 0.942 |
| $\Delta$ | **+0.024** | **+0.009** | **+0.003** | **+0.022** | **-.002** | **+0.005** | **+0.012** | **+0.001** | **+0.003** | **-.001** | **+0.021** |

**Table D1:** Separating co-evolutionary from geometric signal. We compare two independent augmentations of the same pLM encoder: + MSA-PF provides a control for MSA-derived co-evolutionary information, whereas + trunk uses the Protenix-v2 trunk representation, which incorporates both MSA-conditioned and geometry-aware information. Importantly, the trunk is *not* added on top of MSA-PF; both are separately combined with the original pLM baseline. Accordingly, Δ_MSA-PF_ and Δ_trunk_ denote the change in Spearman correlation relative to the same pLM baseline. Positive and negative changes indicate gains and losses, respectively.

| Encoder | Amylase |  |  |  | IRED | Hydro |  |  |  |  |
| --- | --- | --- | --- | --- | --- | --- | --- | --- | --- | --- |
|  | 1→m | c2f | f2c | by-mut |  | 3→m | l2h | P06241 | P0A9X9 | P01053 |
| Sequence-input pLM |  |  |  |  |  |  |  |  |  |  |
| ESM-C | 0.694 | 0.481 | 0.485 | 0.660 | 0.197 | 0.684 | 0.397 | 0.395 | 0.263 | 0.329 |
| ↪ + MSA-PF | 0.728 | 0.610 | 0.548 | 0.691 | 0.195 | 0.677 | 0.333 | 0.399 | 0.280 | 0.177 |
| $\Delta_{\text{MSA-PF}}$ | +0.034 | +0.130 | +0.063 | +0.031 | −0.002 | −0.007 | −0.064 | +0.004 | +0.017 | −0.153 |
| ↪ + trunk | 0.695 | 0.698 | 0.612 | 0.720 | 0.196 | 0.679 | 0.416 | 0.333 | 0.200 | 0.324 |
| $\Delta_{\text{trunk}}$ | +0.001 | +0.217 | +0.127 | +0.060 | −0.001 | −0.006 | +0.020 | −0.062 | −0.063 | −0.005 |
| ESM2 | 0.669 | 0.397 | 0.420 | 0.671 | 0.211 | 0.663 | 0.336 | 0.428 | 0.207 | 0.406 |
| ↪ + MSA-PF | 0.738 | 0.580 | 0.557 | 0.678 | 0.211 | 0.671 | 0.403 | 0.425 | 0.208 | 0.296 |
| $\Delta_{\text{MSA-PF}}$ | +0.069 | +0.183 | +0.137 | +0.007 | .000 | +0.008 | +0.067 | −0.003 | +0.001 | −0.110 |
| ↪ + trunk | 0.685 | 0.665 | 0.614 | 0.701 | 0.181 | 0.659 | 0.368 | 0.275 | 0.321 | 0.413 |
| $\Delta_{\text{trunk}}$ | +0.016 | +0.268 | +0.194 | +0.030 | −0.030 | −0.004 | +0.032 | −0.153 | +0.114 | +0.007 |
| DPLM-2 | 0.681 | 0.393 | 0.367 | 0.638 | 0.200 | 0.562 | 0.377 | 0.263 | 0.103 | 0.419 |
| ↪ + MSA-PF | 0.745 | 0.575 | 0.541 | 0.675 | 0.207 | 0.602 | 0.366 | 0.384 | 0.146 | 0.326 |
| $\Delta_{\text{MSA-PF}}$ | +0.064 | +0.182 | +0.174 | +0.037 | +0.007 | +0.040 | −0.011 | +0.121 | +0.043 | −0.093 |
| ↪ + trunk | 0.700 | 0.661 | 0.593 | 0.704 | 0.175 | 0.583 | 0.350 | 0.229 | 0.048 | 0.247 |
| $\Delta_{\text{trunk}}$ | +0.019 | +0.268 | +0.226 | +0.066 | −0.025 | +0.021 | −0.027 | −0.034 | −0.055 | −0.172 |
| Structure-input pLM |  |  |  |  |  |  |  |  |  |  |
| SaProt | 0.677 | 0.532 | 0.446 | 0.654 | 0.210 | 0.657 | 0.403 | 0.608 | 0.372 | 0.435 |
| ↪ + MSA-PF | 0.737 | 0.601 | 0.548 | 0.658 | 0.213 | 0.665 | 0.428 | 0.562 | 0.280 | 0.370 |
| $\Delta_{\text{MSA-PF}}$ | +0.060 | +0.069 | +0.102 | +0.004 | +0.003 | +0.008 | +0.025 | −0.046 | −0.092 | −0.065 |
| ↪ + trunk | 0.695 | 0.670 | 0.591 | 0.690 | 0.186 | 0.662 | 0.441 | 0.269 | 0.289 | 0.393 |
| $\Delta_{\text{trunk}}$ | +0.018 | +0.138 | +0.145 | +0.036 | −0.024 | +0.005 | +0.038 | −0.339 | −0.083 | −0.042 |
| ESM-3 | 0.739 | 0.623 | 0.528 | 0.688 | 0.211 | 0.674 | 0.411 | 0.478 | 0.345 | 0.397 |
| ↪ + MSA-PF | 0.757 | 0.651 | 0.566 | 0.693 | 0.217 | 0.641 | 0.385 | 0.556 | 0.207 | 0.334 |
| $\Delta_{\text{MSA-PF}}$ | +0.018 | +0.028 | +0.038 | +0.005 | +0.006 | −0.033 | −0.026 | +0.078 | −0.138 | −0.063 |
| ↪ + trunk | 0.733 | 0.712 | 0.605 | 0.702 | 0.179 | 0.658 | 0.397 | 0.287 | 0.311 | 0.349 |
| $\Delta_{\text{trunk}}$ | −0.006 | +0.089 | +0.077 | +0.014 | −0.032 | −0.016 | −0.014 | −0.191 | −0.034 | −0.048 |
| DPLM-2-mm | 0.653 | 0.420 | 0.420 | 0.622 | 0.200 | 0.638 | 0.368 | 0.462 | 0.113 | 0.279 |
| ↪ + MSA-PF | 0.731 | 0.600 | 0.551 | 0.670 | 0.210 | 0.655 | 0.397 | 0.483 | 0.236 | 0.229 |
| $\Delta_{\text{MSA-PF}}$ | +0.078 | +0.180 | +0.131 | +0.048 | +0.010 | +0.017 | +0.029 | +0.021 | +0.123 | −0.050 |
| ↪ + trunk | 0.703 | 0.647 | 0.604 | 0.703 | 0.171 | 0.616 | 0.346 | 0.480 | 0.144 | 0.190 |
| $\Delta_{\text{trunk}}$ | +0.050 | +0.227 | +0.184 | +0.081 | −0.029 | −0.022 | −0.022 | +0.018 | +0.031 | −0.089 |

**Table D2:** Best published baselines in Didi et al. (2026) for each evaluated split. For each split reported in Table 1, we list the strongest published Spearman correlation from Didi et al. (2026) and the corresponding method.

| Dataset | Split | Spearman $\rho$ | Best published method |
| --- | --- | --- | --- |
| Amylase | 1→m | 0.597 | Ridge + likelihood |
|  | c→f | 0.617 | ESM2 likelihood |
|  | f→c | 0.598 | ESM2 likelihood |
|  | by-mutation | 0.681 | Ridge + likelihood |
| IRED | 2→m | 0.211 | Ridge + likelihood |
| Hydro | 3→m | 0.604 | CARP naive |
|  | low→high | 0.308 | Dayhoff |
|  | to-P06241 | 0.426 | Dayhoff |
|  | to-P0A9X9 | 0.151 | Dayhoff |
|  | to-P01053 | 0.329 | ESM-C supervised |
| Rhomax | by-WT | 0.379 | CARP likelihood |
| PDZ3 | 1→2 | 0.512 | CARP supervised |

## References

Yo Akiyama, Zhidian Zhang, Olivia Tang, Rachel Seongeun Kim, Milot Mirdita, Martin Steinegger, and Sergey Ovchinnikov. Expanding the scope of protein language modeling to protein-protein interactions with msa pairformer. Cell, 2026.

Christian B Anfinsen. Principles that govern the folding of protein chains. Science, 181(4096): 223–230, 1973.

David Bednar, Koen Beerens, Eva Sebestova, Jaroslav Bendl, Sagar Khare, Radka Chaloupkova, Zbynek Prokop, Jan Brezovsky, David Baker, and Jiri Damborsky. Fireprot: energy-and evolution-based computational design of thermostable multiple-point mutants. PLoS computational biology, 11(11):e1004556, 2015.

Tristan Bepler and Bonnie Berger. Learning protein sequence embeddings using information from structure. In *International Conference on Learning Representations*, volume 2019, pp. 3162, 2019.

Jun Cheng, Guido Novati, Joshua Pan, Clare Bycroft, Akvilė Z^̌^ emgulytė, Taylor Applebaum, Alexander Pritzel, Lai Hong Wong, Michal Zielinski, Tobias Sargeant, et al. Accurate proteome-wide missense variant effect prediction with alphamissense. Science, 381(6664):eadg7492, 2023.

Christian Dallago, Jody Mou, Kadina E Johnston, Bruce J Wittmann, Nicholas Bhattacharya, Samuel Goldman, Ali Madani, and Kevin K Yang. Flip: Benchmark tasks in fitness landscape inference for proteins. bioRxiv, pp. 2021–11, 2021.

Justas Dauparas, Ivan Anishchenko, Nathaniel Bennett, Hua Bai, Robert J Ragotte, Lukas F Milles, Basile IM Wicky, Alexis Courbet, Rob J de Haas, Neville Bethel, et al. Robust deep learning– based protein sequence design using proteinmpnn. Science, 378(6615):49–56, 2022.

Kieran Didi, Sarah Alamdari, Alex X Lu, Bruce Wittmann, Kadina E Johnston, Ava P Amini, Ali Madani, Maya Czeneszew, Christian Dallago, and Kevin K Yang. Flip2: Expanding protein fitness landscape benchmarks for real-world machine learning applications. bioRxiv, pp. 2026– 02, 2026.

Henry Dieckhaus, Michael Brocidiacono, Nicholas Z Randolph, and Brian Kuhlman. Transfer learning to leverage larger datasets for improved prediction of protein stability changes. Proceedings of the national academy of sciences, 121(6):e2314853121, 2024.

Ken A Dill and Justin L MacCallum. The protein-folding problem, 50 years on. science, 338(6110): 1042–1046, 2012.

Albert Escobedo, Gesa Voigt, Andre J Faure, and Ben Lehner. Genetics, energetics and allostery during a billion years of hydrophobic protein core evolution. bioRxiv, pp. 2024–05, 2024.

Maximilian Gantz, Simon V Mathis, Friederike EH Nintzel, Matthew Penner, Paul J Zurek, Tanja Knaus, Vasilis Tseliou, Elie Patel, Daniel Boros, Friedrich-Maximilian Weberling, et al. Micro-droplet screening rapidly profiles a biocatalyst to enable its ai-assisted engineering. BioRxiv, pp. 2024–04, 2024.

Chao Hou, Haiqing Zhao, and Yufeng Shen. Protein language models trained on biophysical dynamics inform mutation effects. Proceedings of the National Academy of Sciences, 123(4): e2530466123, 2026.

Keiichi Inoue, Masayuki Karasuyama, Ryoko Nakamura, Masae Konno, Daichi Yamada, Kentaro Mannen, Takashi Nagata, Yu Inatsu, Hiromu Yawo, Kei Yura, et al. Exploration of natural red-shifted rhodopsins using a machine learning-based bayesian experimental design. Communications biology, 4(1):362, 2021.

Hyosoon Jang, Hyunjin Seo, Yunhui Jang, Seonghyun Park, and Sungsoo Ahn. Boltz is a strong baseline for atom-level representation learning. arXiv e-prints, pp. arXiv–2602, 2026.

John Jumper, Richard Evans, Alexander Pritzel, Tim Green, Michael Figurnov, Olaf Ronneberger, Kathryn Tunyasuvunakool, Russ Bates, Augustin Z^̌^ídek, Anna Potapenko, et al. Highly accurate protein structure prediction with alphafold. nature, 596(7873):583–589, 2021.

Masayuki Karasuyama, Keiichi Inoue, Ryoko Nakamura, Hideki Kandori, and Ichiro Takeuchi. Understanding colour tuning rules and predicting absorption wavelengths of microbial rhodopsins by data-driven machine-learning approach. Scientific reports, 8(1):15580, 2018.

Diederik P Kingma and Jimmy Ba. Adam: A method for stochastic optimization. arXiv preprint arXiv:1412.6980, 2014.

Mingchen Li, Yang Tan, Xinzhu Ma, Bozitao Zhong, Huiqun Yu, Ziyi Zhou, Wanli Ouyang, Bingxin Zhou, Pan Tan, and Liang Hong. Prosst: Protein language modeling with quantized structure and disentangled attention. Advances in Neural Information Processing Systems, 37:35700–35726, 2024.

Yanghao Li, Naiyan Wang, Jianping Shi, Xiaodi Hou, and Jiaying Liu. Adaptive batch normalization for practical domain adaptation. Pattern Recognition, 80:109–117, 2018.

Zeming Lin, Halil Akin, Roshan Rao, Brian Hie, Zhongkai Zhu, Wenting Lu, Nikita Smetanin, Robert Verkuil, Ori Kabeli, Yaniv Shmueli, et al. Evolutionary-scale prediction of atomic-level protein structure with a language model. Science, 379(6637):1123–1130, 2023.

Amy X Lu, Wilson Yan, Kevin K Yang, Vladimir Gligorijevic, Kyunghyun Cho, Pieter Abbeel, Richard Bonneau, and Nathan C Frey. Tokenized and continuous embedding compressions of protein sequence and structure. Patterns, 6(6), 2025.

M Jehanzeb Mirza, Jakub Micorek, Horst Possegger, and Horst Bischof. The norm must go on: Dynamic unsupervised domain adaptation by normalization. In 2022 IEEE/CVF Conference on Computer Vision and Pattern Recognition (CVPR), pp. 14745–14755. IEEE, 2022.

Pascal Notin, Aaron Kollasch, Daniel Ritter, Lood Van Niekerk, Steffanie Paul, Han Spinner, Nathan Rollins, Ada Shaw, Rose Orenbuch, Ruben Weitzman, et al. Proteingym: Large-scale benchmarks for protein fitness prediction and design. Advances in neural information processing systems, 36: 64331–64379, 2023.

Yeonwoo Park, Brian PH Metzger, and Joseph W Thornton. The simplicity of protein sequence-function relationships. Nature Communications, 15(1):7953, 2024.

Saro Passaro, Gabriele Corso, Jeremy Wohlwend, Mateo Reveiz, Stephan Thaler, Vignesh Ram Somnath, Noah Getz, Tally Portnoi, Julien Roy, Hannes Stark, et al. Boltz-2: Towards accurate and efficient binding affinity prediction. BioRxiv, 2025.

Roshan Rao, Joshua Meier, Tom Sercu, Sergey Ovchinnikov, and Alexander Rives. Transformer protein language models are unsupervised structure learners. In International Conference on Learning Representations, 2021.

Steffen Schneider, Evgenia Rusak, Luisa Eck, Oliver Bringmann, Wieland Brendel, and Matthias Bethge. Improving robustness against common corruptions by covariate shift adaptation. Advances in neural information processing systems, 33:11539–11551, 2020.

Meitar Sela, Jonathan R Church, Igor Schapiro, and Dina Schneidman-Duhovny. Rhomax: computational prediction of rhodopsin absorption maxima using geometric deep learning. Journal of chemical information and modeling, 64(12):4630, 2024.

Yash Semlani, Ruitong Li, Frederick Hoffman, Nauman Javed, and Krithik Ramesh. Isoplm: Isolating the impacts of architecture on protein language models. In The 2026 Workshop on Generative and Agentic AI for Biology, 2026.

Jin Su, Chenchen Han, Yuyang Zhou, Junjie Shan, Xibin Zhou, and Fajie Yuan. Saprot: Protein language modeling with structure-aware vocabulary. In *International Conference on Learning Representations*, volume 2024, pp. 6987–7009, 2024.

Itay Talpir and Sarel J Fleishman. Simple baselines rival protein language models in mutation-dense design tasks. bioRxiv, pp. 2026–05, 2026.

Chai Discovery team, Jacques Boitreaud, Jack Dent, Matthew McPartlon, Joshua Meier, Vinicius Reis, Alex Rogozhonikov, and Kevin Wu. Chai-1: Decoding the molecular interactions of life. BioRxiv, pp. 2024–10, 2024.

Neil Thomas, David Belanger, Chenling Xu, Hanson Lee, Kathleen Hirano, Kosuke Iwai, Vanja Polic, Kendra D Nyberg, Kevin G Hoff, Lucas Frenz, et al. Engineering highly active nuclease enzymes with machine learning and high-throughput screening. Cell systems, 16(3), 2025.

Kotaro Tsuboyama, Justas Dauparas, Jonathan Chen, Elodie Laine, Yasser Mohseni Behbahani, Jonathan J Weinstein, Niall M Mangan, Sergey Ovchinnikov, and Gabriel J Rocklin. Mega-scale experimental analysis of protein folding stability in biology and design. Nature, 620(7973):434–444, 2023.

Floris van der Flier, Dave Estell, Sina Pricelius, Lydia Dankmeyer, Sander van Stigt Thans, Harm Mulder, Rei Otsuka, Frits Goedegebuur, Laurens Lammerts, Diego Staphorst, et al. Enzyme structure correlates with variant effect predictability. 2024.

Michel van Kempen, Stephanie S Kim, Charlotte Tumescheit, Milot Mirdita, Cameron LM Gilchrist, Johannes Söding, and Martin Steinegger. Foldseek: fast and accurate protein structure search. Biorxiv, pp. 2022–02, 2022.

Xinyou Wang, Zaixiang Zheng, Fei Ye, Dongyu Xue, Shujian Huang, and Quanquan Gu. Dplm-2: A multimodal diffusion protein language model. In *International Conference on Learning Representations*, volume 2025, pp. 35463–35490, 2025.

James C Whisstock and Arthur M Lesk. Prediction of protein function from protein sequence and structure. Quarterly reviews of biophysics, 36(3):307–340, 2003.

Talal Widatalla, Ashir A Borah, Samuel H King, Claudia L Driscoll, Rafael Rafailov, and Brian L Hie. Aligning protein-generative models to experimental fitness with proteindpo. Nature Methods, pp. 1–9, 2026.

Phillip R Woolley, Aaron L Feller, Andrew D Ellington, and Claus O Wilke. Overestimating zero-shot fitness prediction: Broad benchmarks mask local failures and practical limitations. bioRxiv, pp. 2026–06, 2026.

Kevin K Yang, Nicolo Fusi, and Alex X Lu. Convolutions are competitive with transformers for protein sequence pretraining. Cell systems, 15(3):286–294, 2024.

Kevin K Yang, Sarah Alamdari, Alex J Lee, Kaeli Kaymak-Loveless, Samir Char, Garyk Brixi, Carles Domingo-Enrich, Chentong Wang, Suyue Lyu, Nicolo Fusi, et al. The dayhoff atlas: scaling sequence diversity for improved protein generation. bioRxiv, pp. 2025–07, 2025.

Taraneh Zarin and Ben Lehner. A complete map of specificity encoding for a partially fuzzy protein interaction. BioRxiv, 2024.

Yuxuan Zhang, Chengyue Gong, Jinyuan Sun, Jiaqi Guan, Milong Ren, Song Xue, Hanyu Zhang, Wenzhi Ma, Zhenyu Liu, Xinshi Chen, et al. Protenix-v2: Broadening the reach of structure prediction and biomolecular design. bioRxiv, pp. 2026–04, 2026.

